# Coordinated regulation of ribosomes and proteasomes by PRMT1 in the maintenance of neural stemness of cancer cells and neural stem cells

**DOI:** 10.1101/2020.12.17.423362

**Authors:** Lu Chen, Min Zhang, Lei Fang, Xiaoli Yang, Liyang Xu, Lihua Shi, Ning Cao, Ying Cao

## Abstract

Our studies suggest that neural stemness contributes to cell tumorigenicity. The basic cell physiological machineries and developmental programs, such as cell cycle, ribosomes, proteasomes, epigenetic factors, etc., which are upregulated in and promote cancers, are enriched in embryonic neural cells. How these machineries are coordinated is unknown. Here, we show that loss of neural stemness in cancer cells or neural stem cells leads to simultaneous downregulation of components of ribosomes and proteasomes, which are responsible for protein synthesis and degradation, respectively, and downregulation of major epigenetic factors. Inhibition of PRMT1 causes neuron-like differentiation, downregulation of a similar set of proteins, and alteration of subcellular localization of ribosome and proteasome components. PRMT1 interacts with these components, catalyzes arginine methylation of them and protects them from degradation, thereby maintaining a high level of expression of epigenetic factors that maintain neural stemness. PRMT1 inhibition results in repression of cell tumorigenicity. Therefore, PRMT1 coordinates ribosomes and proteasomes to match the needs for high protein production and protein homeostasis in cells with fast cell cycle and proliferation.

## Introduction

Ribosomes and proteasomes are machineries that are required for basic cellular physiology. Ribosomes are responsible for protein translation from mRNAs. In eukaryotic cells, the mature 80S ribosome is composed of the 40S and 60S subunits. In human, the 40S subunit contains 18S ribosomal RNA (rRNA) and 33 ribosomal proteins, such as RPS2, RPS3, RPS3A, RPS4X, RPS5-RPS21, and RPS23-RPS29, etc.; while the 60S subunits contains 5S, 5.8S and 28S rRNAs and 47 ribosomal proteins, mainly RPL3-RPL5, RPL7-RPL19, RPL21-RPL24, RPL23A, RPL26-RPL31, etc. (de la Cruz et al., 2015; Pelletier et al., 2018). Some additional proteins are assembled into pre-40S and pre-60S ribosomal subunits in the nucleoplasm and cytoplasm or required for nuclear export of the pre-60S subunit. These include the nuclear chaperone MDN1 (Midasin), an ATPase critical for proper remodeling of pre-60S subunit (Konikkat and Woolford, 2017; Kressler et al., 2012). The major steps of ribosome biogenesis occur in nucleolus. At late stage of maturation, 80S ribosomes are exported to cytoplasm, where they undergo the last step of maturation and start protein synthesis (Pelletier et al., 2018). Therefore, ribosomes are essential for cellular survival, growth and differentiation. An increase in overall protein levels and production of sufficient ribosomes are a prominent characteristic of cell growth (Baserga, 2007; Turi et al., 2019). Ribosome biogenesis is indispensible for maintaining pluripotency (Watanabe-Susaki et al., 2014). Vice versa, ribosomes exhibit the ability to transdifferentiate human somatic cells into multipotent state (Ito et al., 2018a, b). These imply the critical roles of ribosomes in early development and cell differentiation.

While protein synthesis is fundamental for normal physiology of a cell, the reverse process, protein degradation is equally important, because unneeded or damaged proteins are detrimental to a cell. Proteasomes are multi-enzyme complexes responsible for degrading excessive or wrong proteins, in cooperation with ubiquitin. The 26S proteasome (or proteasome) is the major cellular protease, found in the nucleus and cytoplasm of all eukaryotic cells and playing the central role in the ubiquitin-dependent pathway of protein degradation (Konstantinova et al., 2008; Bedford et al., 2010; Schröter and Adjaye, 2014). The proteasome consists of a 20S core particle that is responsible for catalytic activity and two 19S regulatory particles that recognize ubiquitinated and misfolded proteins. The 20S catalytic core is composed of two α-rings and two β-rings. A α-ring is made up of seven components PSMA1-PSMA7, while a β-ring includes PSMB1-PSMB7. 19S regulatory particle has more complex constituents, including PSMC1-PSMC6, PSMD1-PSMD4, PSMD6-PSMD8, PSMD11-PSMD14, and ADRM1 (Konstantinova et al., 2008; Bedford et al., 2010; Schröter and Adjaye, 2014; Rousseau and Bertolotti, 2018). Other regulatory complexes such as the PA28 activator, consisting of PSME1-PSME3, associate with and activate the 20S core (Konstantinova et al., 2008; Schröter and Adjaye, 2014). Proteasomes are fundamental to life process because of its central function in maintaining protein homeostasis, a status essential for cell development and relieving cellular ageing (Bedford et al., 2010). Particularly, proteasomes play a key role in promoting self-renewal of neural progenitor cells (NPCs) (Zhao et al., 2016)

Protein arginine methylation is a common type of post-translational modification that is catalyzed by either one of the nine protein arginine methyltransferases (PRMT1-PRMT9). PRMT1 is the main epigenetic factor responsible for asymmetrical dimethylation histone H4 at Arg-3 and functions as a transcriptional co-activator. Besides, PRMT1 also methylates nonhistone proteins, thereby modulating many biological processes such as RNA metabolism, genome stability, transcription, and signal transduction (Blanc and Richard, 2017; Yang and Bedford, 2013). Mouse embryos without Prmt1 cannot develop beyond E6.5 (Pawlak et al., 2000). Other major epigenetic factors like EZH2, LSD1, DNMT1, HDAC1, etc., which mediate different types of epigenetic modifications, are essential for embryonic developmental programs. Loss of either of them causes early embryonic lethality. They regulate neuronal differentiation from neural stem/progenitor cells (NSCs/NPCs) through different mechanisms (Gaub et al., 2010; Cho and Cavalii, 2014; Han et al., 2014; Lei et al., 2019).

Tumorigenesis appears as an enormously complex process with dysregulation of numerous regulatory signals and networks. However, cancer cells (tumor-initiating cells) are characteristic of NSCs/NPCs because they have neuronal differentiation potential and most cancer promoting genes or genes upregulated in cancer are embryonic neural specific, indicating a similarity of regulatory networks between cancer cells and NSCs/NPCs (Zhang et al., 2017; Cao, 2017; Lei et al., 2019). More recently, we demonstrated that cell tumorigenicity and differentiation potential stem from neural stemness, a property that is pre-determined by evolutionary advantage (Xu et al., 2020). Together with the evidence from studies on tumor biology and developmental biology, we propose that neural stemness represents the ground or basal state of cell tumorigenicity and differentiation potential (Cao, 2020). Cancer cells are characteristic of fast cell cycle and proliferation, a feature that needs high level of protein production. In agreement, ribosome biogenesis is upregulated in cancer cells and involved in cancer initiation and progression (Bustelo and Dosil, 2018; Catez et al., 2018; Pelletier et al., 2018; Turi et al., 2019). Enhanced protein synthesis means a higher demand for regulatory machinery maintaining protein homeostasis. Actually, cancer cells show elevated levels of proteasomes and proteasome activity for protein quality control to promote their survival, growth and metastasis (Chen et al., 2017; Mofers et al., 2017; Soave et al., 2017; Rousseau and Bertolotti, 2018). Therefore, a concerted regulation of ribosome and proteasome is required for tumorigenesis. PRMT1 and a series of epigenetic factors, such as EZH2, LSD1, DNMT1 or HDAC1, are also upregulated in various cancer types and promote cancer (Yang and Bedford, 2013; Baldwin et al., 2014; Bennett and Licht, 2018; Mohammad et al., 2019). Consistent with the dysregulated expression and roles of these proteins during tumorigenesis is the specific or enriched expression of the genes for ribosome and proteasome components and epigenetic factors in embryonic neural cells or NSCs during vertebrate embryogenesis (Cao, 2020). Here, we show the evidence that coordinated regulation of ribosome biogenesis and proteasome by PRMT1 is required for maintenance of neural stemness in both cancer cells and NSCs, which are highly proliferative and need a high protein production and protein homeostasis.

## Results

### Loss of PRMT1 function leads to neuronal differentiation in both cancer cells and neural stem cells

PRMT1 functions as an oncoprotein. Correspondingly, its gene shows strongly enriched expression in neural precursor tissues and developing central nervous system during early vertebrate embryogenesis (Pawlak et al., 2000; Zhang et al., 2017), suggesting that it might play a role in maintaining neural stemness. When PRMT1 was knocked down using a validated hairpin RNA (shPRMT1) in lung cancer cell line A549, the cells showed neuron-like phenotype (Figure 1A) with extended neuritic processes. Immunofluorescence (IF) confirmed a successful knockdown of PRMT1 (Figure 1B). A key pro-neuronal differentiation protein NEUROD1 and neuronal markers TUBB3 and MAP2 were not detected in control cells (shCtrl), but detected in knockdown cells (Figure 1B). This phenotypic change was recapitulated in melanoma cell line A375 after knockdown of PRMT1, as shown by neurite-like extensions in knockdown cells (Figure 1C). IF demonstrated efficient knockdown effect of shPRMT1 and expression of neuronal markers TUBB3 and NF-L in the knockdown cells (Figure 1D). We then tested the effect of Prmt1 on differentiation of primitive neural stem cells (primNSCs) derived from mouse embryonic stem cells (mESCs). Knockdown of Prmt1 in mESCs led to formation of clones of reduced size as compared with control cells on feeder cells in ESC medium, but no significant differentiation effect was observed (Figure S1). When cultured in NSC-specific serum-free medium, however, control cells formed free-floating neurospheres (Figure 1E), which grew larger gradually in an extended culture period. In contrast, knockdown cells formed clusters attached to the bottom of culture dish, showing a phenotype of neuronal differentiation (Figure 1E). IF detection revealed expression of Prmt1, the neural stemness markers Sox1 and Pax3 in neurospheres (Figure 1F). Prmt1 was effectively inhibited by shPrmt1. Meanwhile, neural stemness markers were lost in the knockdown cells (Figure 1F). Neuronal proteins Neurod1, Tubb3 and Map2 were not present in neurospheres. Prmt1 knockdown resulted in expression of these proteins in cells (Figure 1F), indicating neuronal differentiation of primNSCs in response to Prmt1 inhibition. Loss of PRMT1/Prmt1 function leads to neuronal-like differentiation in cancer cells and NSCs, an effect similar to our previous observations on inhibition of other cancer-promoting factors in cancer cells and NSCs (Zhang et al., 2017; Lei et al., 2019). This suggests that PRMT1/Prmt1 functions in conferring or maintaining neural stemness in both cancer cells and NSCs.

**Figure 1.**
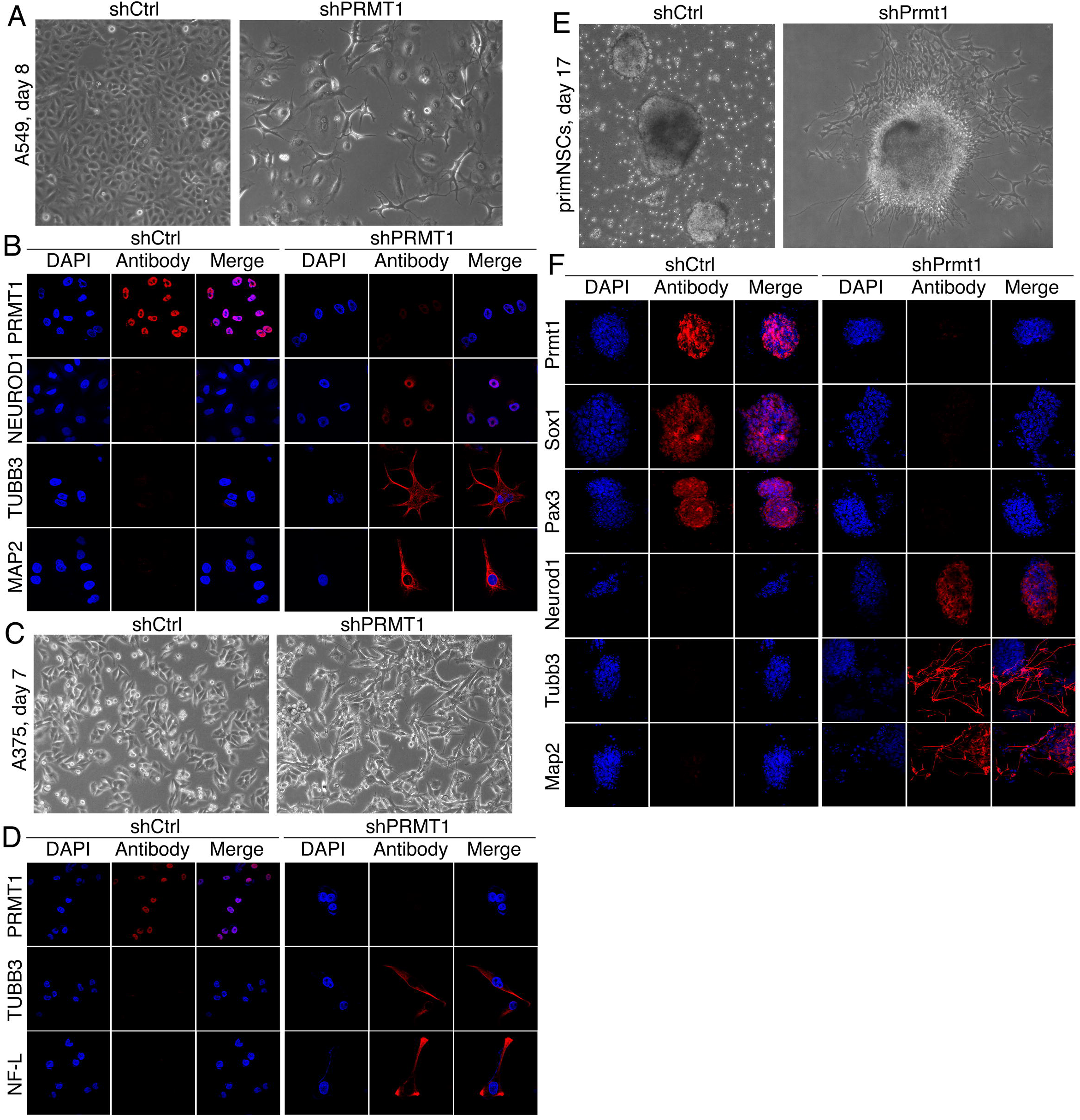
Neuronal differentiation effect induced by knockdown of PRMT1/Prmt1 in cancer cells or NSCs. (A, B) Neuronal-like differentiation phenotype by knockdown of PRMT1 (shPRMT1) in A549 cells in a period of 8 days (A), and validation of knockdown effect of PRMT1 and detection of neuronal protein expression (NEUROD1, TUBB3, MAP2) in cells using immunofluorescence (IF) (B). Cells infected with lentivirus containing empty vector (shCtrl) were used as control. (C, D) Neuronal-like differentiation phenotype by knockdown of PRMT1 in A375 cells in a period of 7 days (A), and validation of knockdown effect of PRMT1 and detection of neuronal marker expression (TUBB3, NF-L) in cells using IF (B). (E) Neurosphere formation of primNSCs derived from mouse ESCs in NSC serum-free medium (left) and neuronal differentiation phenotype induced by knockdown of Prmt1 in primNSCs (right). (F) Analysis of Prmt1 knockdown effect and detection of neural stemness markers (Sox1, Pax3) and neuronal proteins (Neurod1, Tubb3, Map2) in neurospheres and in cell clusters with Prmt1 knockdown using IF. In all IF assays, nuclei were counterstained with DAPI.

### Loss of neural stemness via differentiation causes downregulation of basic cellular machineries, and PRMT1 regulates ribosomes and proteasomes

To explore how PRMT1/Prmt1 plays its functions in cancer cells and NSCs, we tried to identify potential PRMT1/Prmt1 interaction proteins using mass spectrometry. In NE-4C cells, an NSC cell line that was derived from cerebral vesicles of mouse E9 embryos, 1629 proteins were identified as putative Prmt1 interaction partners (Table S1). In hepatocellular carcinoma cell line HepG2, 2020 proteins were identified (Table S2). There are 1083 putative interaction partners in common (Figure 2A; Tables S1 and S2), suggesting that PRMT1/Prmt1 mediates largely similar regulatory networks in both cell types. The 1629 proteins in NE-4C cells are mostly enriched in pathways that underlie basic cellular physiological functions, such as spliceosome, RNA transport, ribosome, proteasome, etc. (Figure 2B). In agreement, the most enriched gene ontology (GO) terms for molecular functions are poly(A) RNA and RNA binding, etc, which are associated biological processes of translation, mRNA processing, RNA splicing, etc., that should occur in nucleus and cytoplasm (Figure 2B). The 2020 proteins in HepG2 cells are mainly enriched in similar pathway and GO terms (Figure 2C). Noteworthy is that the enriched terms for PRMT1/Prmt1 interaction proteins are associated with basic cellular physiological functions.

**Figure 2.**
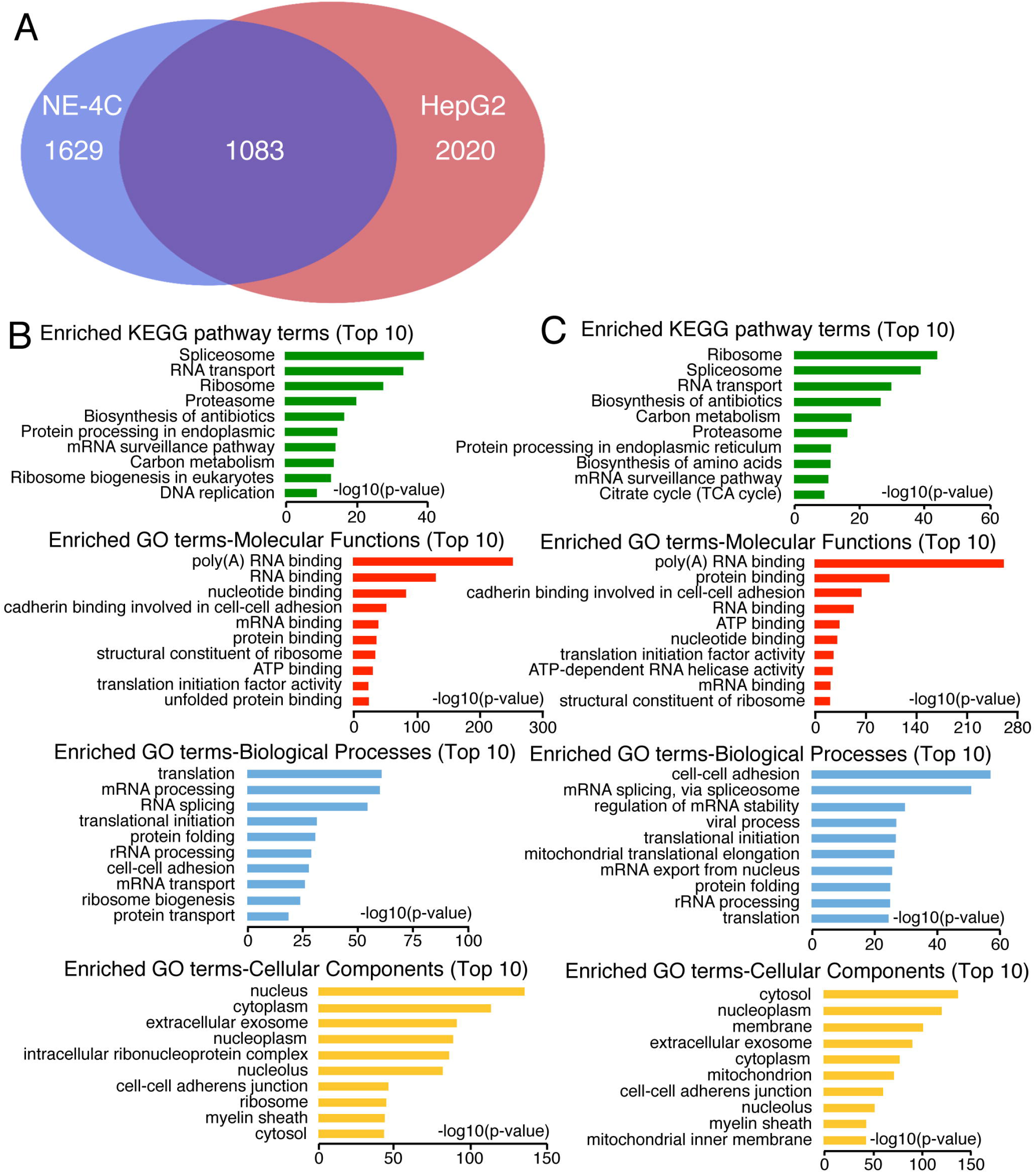
Identification of putative interaction proteins of PRMT1/Prmt1 using mass spectrometry, and bioinformatic analysis on the identified proteins. (A) A diagram summarizing the number of identified interaction partners of Prmt1/PRMT1 in NE-4C cells and HepG2 cells, and the shared interaction partners between the two cells. (B, C) Bioinformatic analysis on enriched pathway and GO terms in the interaction partners in NE-4C (B) or HepG2 (C) cells.

Many ribosome and proteasome component proteins were identified as PRMT1/Prmt1 putative interaction partners in the mass spectrometric assays (Tables S1 and S2). Intriguingly, expression of the genes for these components, as well as the genes for typical epigenetic factors, such as PRMT1, HDAC1, DNMT1, SETDB1, EZH2 and LSD1, is enriched in embryonic neural cells during vertebrate embryogenesis (Figure S2), suggesting that they work together to regulate neural development. We examined first how these proteins are regulated during neuronal differentiation. Treatment with retinoic acid (RA), an agent inducing neuronal differentiation from NSCs, caused neuronal differentiation in NE-4C cells (Figure 3A). The differentiation led to a decreased expression in proteasome components such as Psma2, Psmd2, Psmd4 and Adrm1, and ribosome components such as Rps2, Rps3, Rpl24 and Rpl26 (Figure 3B). Decreased proteins also included epigenetic modification factors Ezh2, Setdb1 and Lsd1 in addition to Prmt1. These are oncoproteins, and Ezh2 and Lsd1 are known to maintain neural stemness (Zhang et al., 2017; Lei et al., 2019; Han et al., 2014). Neural stemness proteins Zic1, Msi1 and Pax6 were decreased and neuronal proteins Nf-l, Tubb3, Map2 and Syn1 were correspondingly upregulated (Figure 3B). We also examined the effect of neuronal differentiation induced by PRMT1/Prmt1 knockdown on the expression of these proteins. In NE-4C cells, knockdown of Prmt1 also generated a neuronal phenotype (Figure 3C). The neural stemness marker Pax6 was downregulated and the neuronal protein Map2 was upregulated in knockdown cells (Figure 3D). Likewise, the proteasome and ribosome components and epigenetic factors were simultaneously downregulated (Figure 3D). IF also demonstrated that neural stemness markers Sox1 and Pax6, which were expressed in control NE-4C cells, were significantly reduced in cells with knockdown of Prmt1 (Figure S3A). In contrast, the neuronal markers, Nf-l and Tubb3, which were not expressed in control cells, were expressed in long neurite-like processes in knockdown cells (Figure S3B), supporting again the neuronal differentiation effect. The similar tendency of downregulation of these proteins was observed in A549 cells after PRMT1 knockdown (Figure 3E). Besides, PCNA, AURKA, SOX1/2 and MYC, which promote tumorigenesis, neural stemness or cell cycle, were reduced. Meanwhile, neuronal proteins NEUROD1, NF-L and TUBB3 were increased (Figure 3E). The mode of expression change of these proteins in in vitro neuronal differentiation resembles what occurs during normal neural development. With the progression of neuronal differentiation in mouse embryos from E13.5 to E15.5, epigenetic factors, the ribosome and proteasome proteins and cell cycle protein in embryonic cortical cells are significantly reduced, accompanied with an increase in neuronal proteins (Figure 3F). Moreover, downregulation of ribosome biogenesis during early forebrain development is reported (Chau et al., 2018). In addition to reduced expression levels of ribosome and proteasome proteins, their cellular localization changes in response to loss of PRMT1. In control A375 cells, PRS3 displayed a ubiquitous distribution in a cell with strong enrichment in nucleoli, the site of ribosome biogenesis. By contrast, it was mainly distributed outside nucleus in cells with PRMT1 knockdown (Figure 3G). In control cells, RPL26 was almost uniformly distributed in both nuclei and cytoplasm except its absence in nucleoli. Nevertheless, it was detected primarily outside the nuclei in knockdown cells (Figure 3G). Proteasome protein PSMD2 is also uniformly expressed in both nuclei and cytoplasm with the absence in nucleoli. Similarly, PRMT1 knockdown caused an exclusively cytoplasmic distribution of PSMD2 (Figure 3G). The 20S proteasomes, which can be recognized by the proteasome 20S α+β antibody (20S α+β), were primarily detected in the nuclei of control cells. In knockdown cells, their distribution was nearly uniform throughout the cells (Figure 3G). Contrary to the effect of knockdown, overexpression of PRMT1 was able to enhance strongly the expression of RPS3 in the nucleoli and the expression of PSMD2 in cell nuclei (Figure 3H).

**Figure 3.**
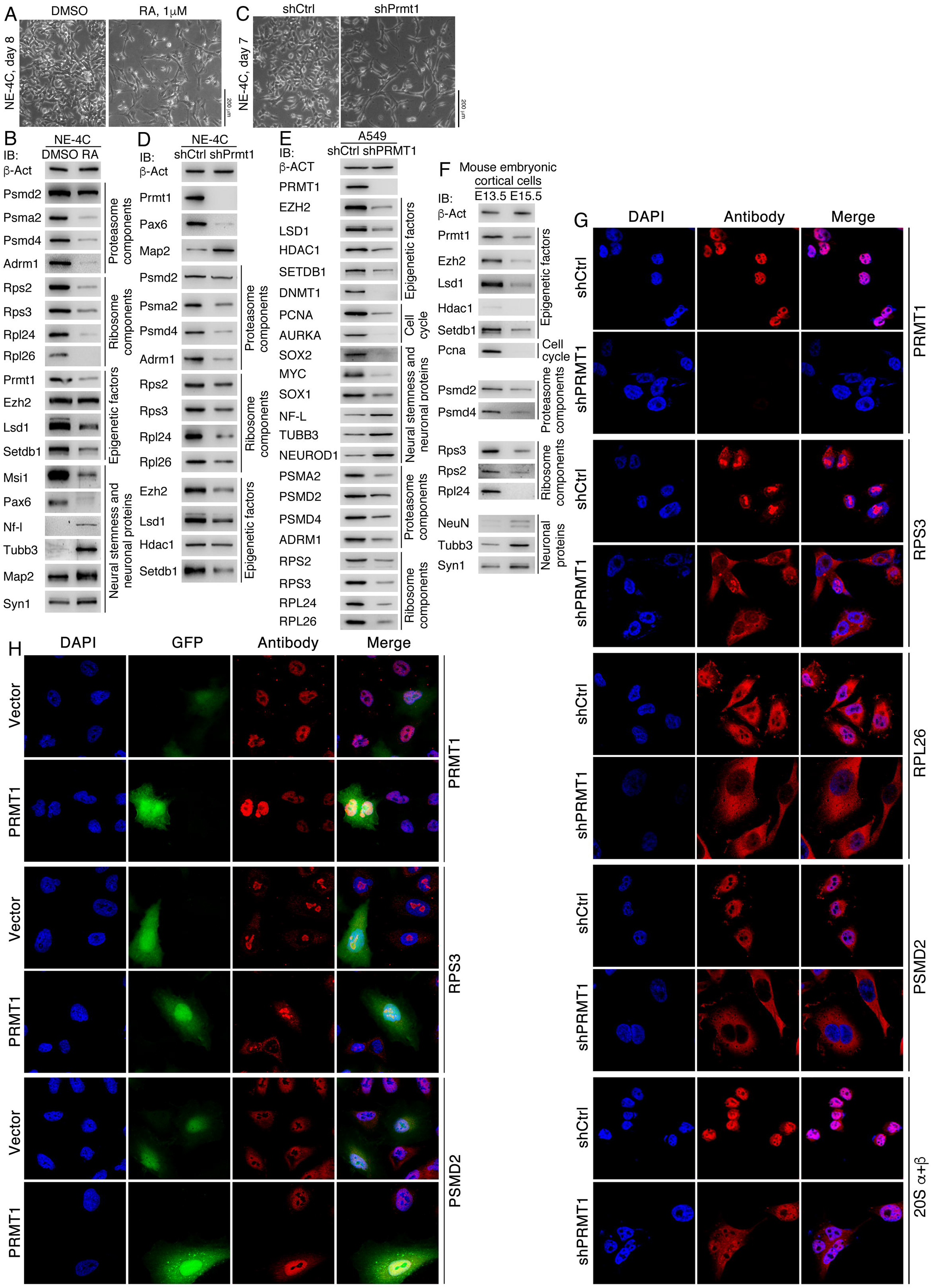
Coordinated regulation of ribosome and proteasome component proteins during neuronal differentiation. (A, B) RA induced neuronal differentiation phenotype in NE-4C cells shown at the time indicated after treatment (A), and immunoblotting (IB) detection of a series of proteasome, ribosome components, epigenetic factors, and neural stemness and neuronal proteins in control cells treated with vehicle (DMSO) and cells treated with RA, as indicated (B). (C, D) Neuronal differentiation phenotype in NE-4C cells induced by Prmt1 knockdown at the indicated time (C), and IB detection of expression of proteins, as indicated, in control (shCtrl) and knockdown (shPrmt1) cells (D). (E) IB detection of various proteins, as indicated, in control A549 (shCtrl) cells and cells with knockdown of PRMT1 (shPRMT1). (F) IB detection of various proteins, as indicated, in mouse embryonic cortical cells at two developmental stages. (G) IF detection of the effect of PRMT1 knockdown on the subcellular distribution of ribosome components RPS3 and RPL26, proteasome protein PSMD2 and 20S subunit (20S α+β) in A375 cells. In (B), (D), (E) and (F), whole cell lysates were used for IB, and detection of β-Act was used as a loading control. (H) IF detection of the effect of PRMT1 overexpression on the expression of ribosome and proteasome proteins in A549 cells. Cells infected with the virus containing only the vector (Vector) as a control.

MYOD1 is a key factor driving myogenesis and can induce muscle cell-like differentiation in different cancer cells (Weintraub et al., 1989). Forced expression of MYOD1 led to the gain of prominent muscle cell phenotype in A375 cells (Figure 4A), concurrent with activation of the muscle cell marker MEF2C, and downregulation of proteasome component proteins PSMA2, PSMD2, PSMD4 and ADRM1, ribosome component proteins RPS2, RPS3, RPL24 and RPL26, and a series of epigenetic factors DNMT1, SETDB1, HDAC1, LSD1, PRMT1 and EZH2 (Figure 4B). Gain of muscle cell-like phenotype and similar pattern of protein expression change were also observed in A549 cells with forced MYOD1 expression (Figure 4C, D). IF assays showed activation of MEF2C expression in response to MYOD1 expression. Expression of GFP (ZsGreen) alone did not cause any significant change (Figure 4E). In control cells expressing GFP, RPS3 was enriched in nucleoli, and GFP did not affect this distribution. However, expression of MYOD1 caused RPS3 to distribute dominantly in cytoplasm, accompanied with the loss of enrichment in nucleoli (Figure 4E). RPL26 was expressed in both nuclei and cytoplasm in control cells; whereas in cells expressing MYOD1, the nuclear fraction of RPL26 expression was reduced, rendering the protein to distribute primarily in cytoplasm. Proteasome component PSMA2 showed expression in both nuclei and cytoplasm, with slightly higher expression in nuclei; MYOD1 caused a reduction of its nuclear expression (Figure 4E). The proteasome 20S subunit (20S α+β) displayed a similar pattern of expression in control cells. In cells expressing MYOD1, it was expressed almost entirely in cytoplasm, with a loss of expression in nuclei (Figure 4E). Therefore, muscle cell differentiation driven by MYOD1 expression led to not only the decrease in overall expression level of ribosome and proteasome components, but also their subcellular distribution, an effect similar to what was observed in neuronal differentiation. Recently, we showed that the myoblast C2C12 cells gained the phenotype of NSCs and hence tumorigenicity when the gene for Myod1 was knocked out (Xu et al., 2020). In the present study, we found that the above-mentioned epigenetic factors, proteasome and ribosome component proteins were upregulated in knockout (KO) cells, as compared with wild type (WT) C2C12 cells (Figure 4F). Taken together, reduced neural stemness in both NSCs and cancer cells via differentiation results in reduced expression and altered subcellular distribution of ribosome and proteasome components and a series of epigenetic factors in a coordinated manner. Vice versa, gain of neural stemness leads to an opposite change. Therefore, enrichment of basic machineries is an intrinsic feature of neural stemness, and PRMT1 might play a role in the coordination of these basic machineries.

**Figure 4.**
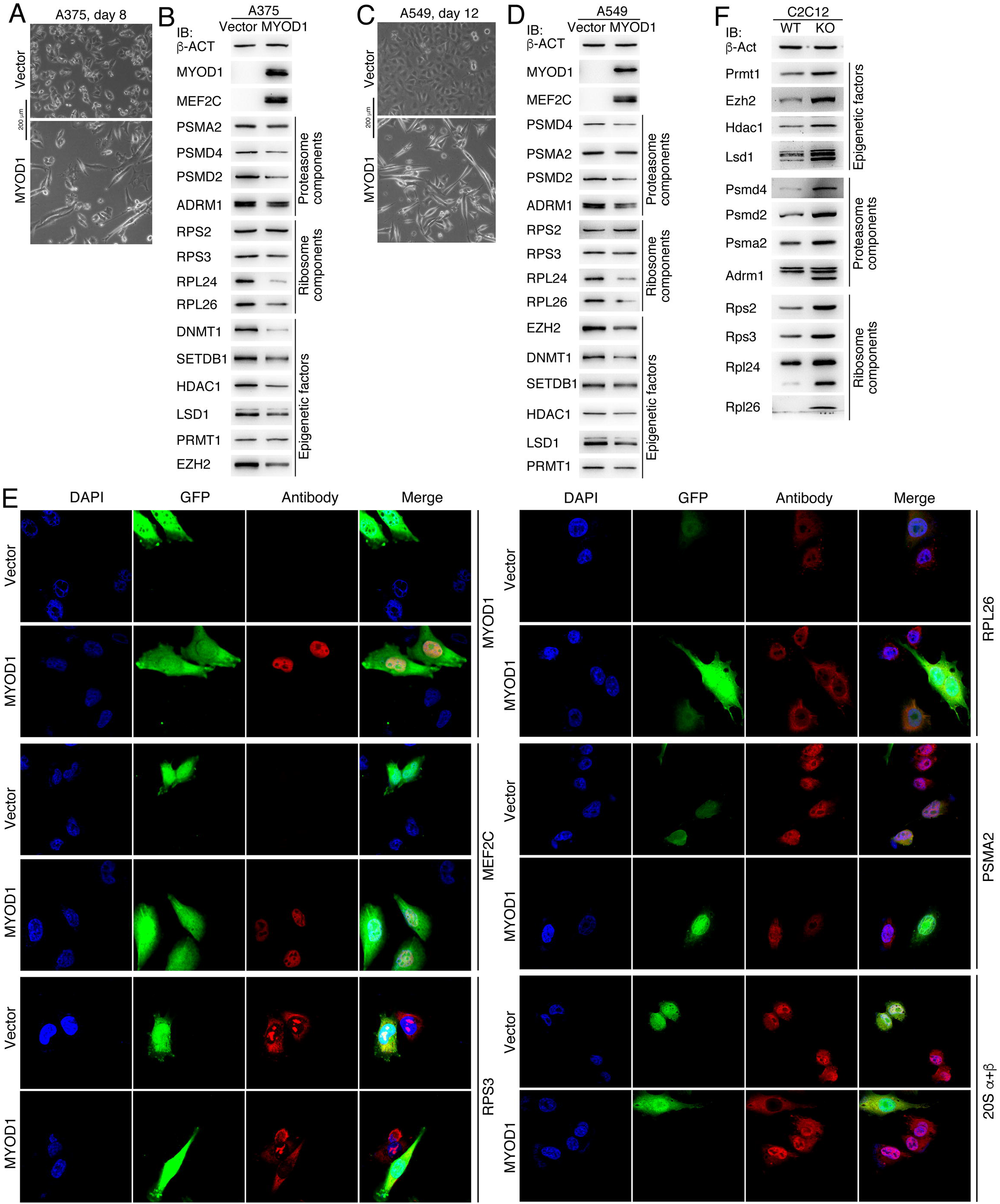
Effect of MYOD1 induced differentiation on the expression of ribosome and proteasome components and epigenetic factors. (A) Phenotypic change in A375 cells infected with virus containing MYOD1. Cells infected with virus containing the empty vector were used as a control. (B) IB detection of expression change of a series of proteins, as indicated, in control A375 cells and cells with forced MYOD1 expression. (C) Phenotypic change in A549 cells in response to MYOD1 expression. (D) Detection of protein expression, as indicated, in control cells and cells with MYOD1 expression using IB. (E) IF detection of expression and subcellular distribution of a muscle cell marker, ribosome and proteasome proteins and the proteasome 20S subunit in control cells and cells with MYOD1 expression. (F) Expression alteration in a series of proteins, as indicated, in wild type (WT) C2C12 cells and the cells with knockout of *Myod1* gene (KO).

### PRMT1 methylates ribosomal and proteasomal proteins, and mediates their proteasomal degradation

Mass spectrometric assays identified ribosome and proteasome component proteins and epigenetic factors as potential interaction partners of PRMT1/Prmt1 (Tables S1 and S2). Protein co-immunoprecipitation (Co-IP) confirmed that PRMT1 interacted with EZH2, LSD1, DNMT1, HDAC1 and SETDB1, the proteasome components PSMD2, PSMD4, PSMA2 and ADRM1, the ribosome components RPS3, RPL24 and RPL26, and with the neural stemness factor MSI1 (Figure 5A). In A375 cells with PRMT1 knockdown, gene expression for epigenetic factors except DNMT1, ribosome and proteasome component proteins were mostly not significantly affected (Figure S4A). Nevertheless, there was a general tendency of upregulation of neuronal genes and downregulation of neural stemness genes, in agreement with neuronal-like differentiation phenotype. Similar patterns of gene expression change were found in A549 cells after PRMT1 knockdown (Figure S4B). Therefore, PRMT1 should regulate the protein expression at post-transcriptional level. We were able to detect arginine methylation on overexpressed RPS3, RPL26 and PSMD2 in HEK293T cells using an antibody against mono- and di-methyl Arginine (Me-Arg) (Figure 5B). PRMT1 knockdown decreased the level of methylated proteins (Figure 5B). By contrast, methylation of these proteins was enhanced when PRMT1 was simultaneously overexpressed (Figure 5C). This suggests that the detected arginine methylation is specifically catalyzed by PRMT1. We observed that inhibition of PRMT1 resulted in an enhanced ubiquitination of these proteins, ultimately leading to their degradation (Figure 5D). Moreover, EZH2, LSD1 and HDAC1 showed an accelerated degradation in cells with PRMT1 inhibition, as compared with the proteins in control cells in a time-course analysis (Figure 5E, F), and also showed an enhanced ubiquitination in response to PRMT1 inhibition (Figure 5G, H). In contrast, expression of EZH2, LSD1 and HDAC1 was enhanced in cells overexpressing PRMT1, as revealed by IF detection (Figure 5I).

**Figure 5.**
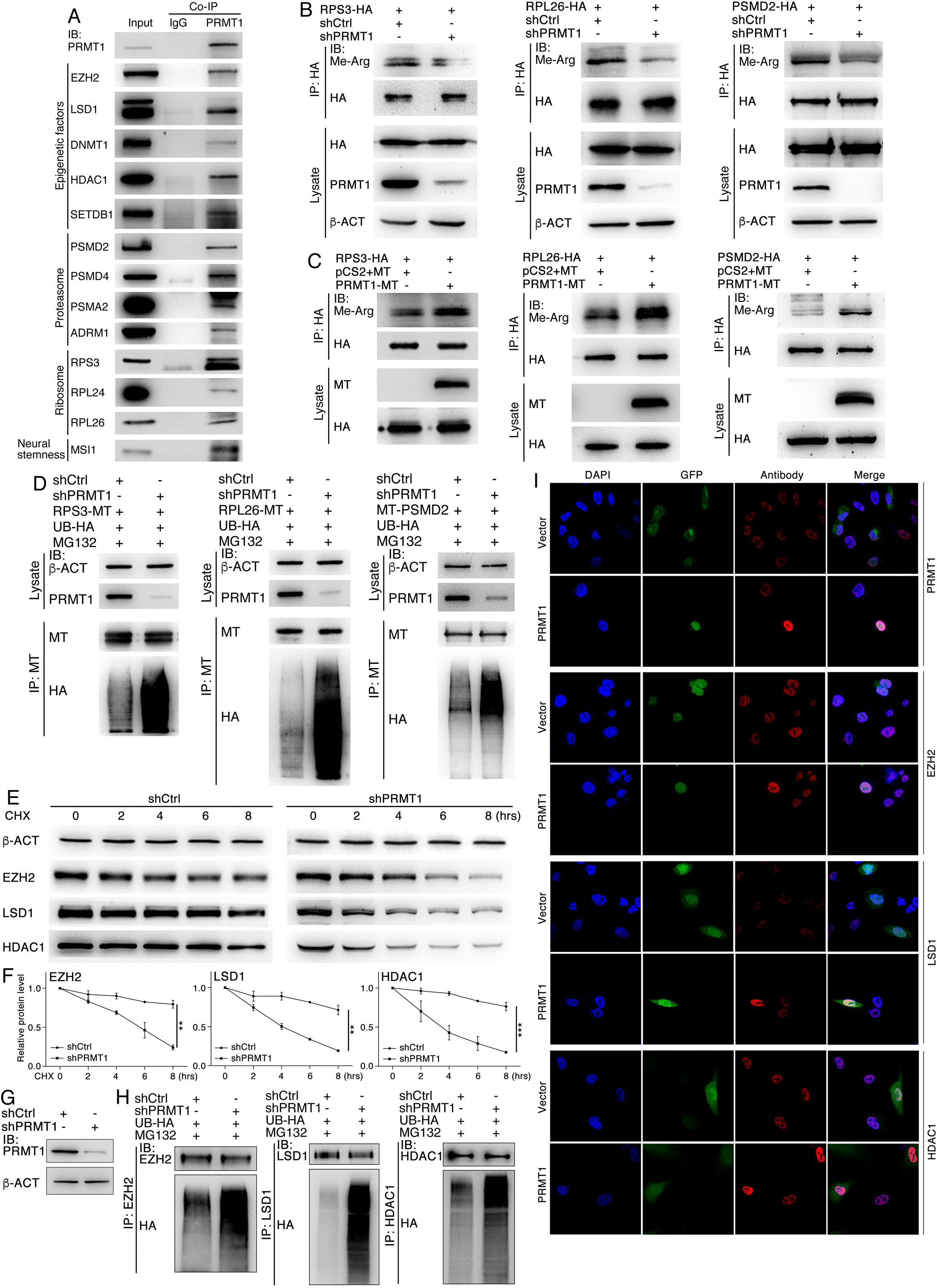
Regulation of ribosome, proteasome component proteins and epigenetic factors by PRMT1. (A) Co-IP confirmation of interaction between PRMT1 and a series of epigenetic factors, ribosome and proteasome proteins and a neural stemness protein, which were identified in mass spectrometry. Immunoprecipitates using IgG were used as negative controls, and proteins in whole cell lysate were used as input. (B) Detection of arginine methylation (Me-Arg) of overexpressed RPS3, RPL26 and PSMD2, and the influence of PRMT1 knockdown (shPRMT1) on arginine methylation of these proteins. (C) The influence of overexpression of PRMT1 on arginine methylation of the proteins. (D) Ubiquitination of overexpressed RPS3, RPL26 and PSMD2, and the effect of PRMT1 knockdown on the ubiquitination of these proteins. (E) The effect of PRMT1 knockdown on protein half-lives by treating cells with CHX in a time series (E). (F) Quantification of relative protein levels in triplicate experiments in (E). Significance was calculated using two-tailed Student’s *t*-test. Data are shown as mean±SEM. **p<0.01, ***p<0.001. (G, H) Influence of PRMT1 knockdown on the ubiquitination of EZH2, LSD1 and HDAC1 (H). (G) shows knockdown effect in cells for ubiquitination assays.

PRMT1 knockdown inhibits both ribosome biogenesis and proteasome assembly, thereby disrupting their functions. We further tested the effect of specific disruption of ribosomal function on the expression of the proteins. MDN1 is required for maturation and nuclear export of pre-60S ribosome subunit (Raman et al., 2016). In cells overexpressing PRMT1, the epigenetic factors, proteasome and ribosome component proteins were upregulated (Figure S5A). When MDN1 was knocked down, most of these proteins were downregulated. If PRMT1 was added to cells with MDN1 inhibition, expression level of the downregulated proteins was increased again (Figure S5A), implying that PRMT1 indeed coordinates expression of the proteins for ribosome biogenesis, proteasome assembly and other epigenetic factors. Disrupted proteasome assembly resulting from PRMT1 inhibition does not mean a total loss of proteasomal activity. In cells treated with either the vehicle (DMSO) or the proteasome inhibitor MG132, PRMT1 knockdown caused downregulation of EZH2, LSD1 and HDAC1 in both cases (Figure S5B). However, MG132 treatment enhanced protein expression in either control or knockdown cells compared with vehicle treated cells, respectively (Figure S5B). In summary, PRMT1 maintains both ribosome biogenesis and proteasome assembly via arginine methylation of their components proteins in NSCs and cancer cells, thus allowing a high capability of protein production and maintenance of protein homeostasis. Lowering the level of PRMT1 causes differentiation of cells and meanwhile decreases both the level of ribosome and proteasome, leading to a lower protein synthesis and correspondingly lower requirement for proteasome activity.

### Inhibition of PRMT1 inhibits malignant features and tumorigenicity of cancer cells and NSCs

Since the epigenetic factors, ribosome biogenesis and proteasome play cancer-promoting roles, their inhibition should generate a repression effect on malignant features and cell tumorigenicity. Fast cell cycle and proliferation is a typical feature of cancer cells. A transcriptomic analysis demonstrated that PRMT1 knockdown in A549 cells led to a general transcriptional change that is mostly associated with cell cycle, mitotic nuclear division, chromosome, protein binding, etc., according to GO, and with cell cycle, DNA replication according to pathway classifications (Figure S6A). This is coherent with the neuronal-like differentiation effect induced by inhibition of PRMT1 (Figure 1A). Furthermore, PRMT1 knockdown strongly inhibited the capability of invasion and migration in A549, A375 and NE-4C cells (Figure S6B, E, H). Meanwhile, compared with control cells, knockdown cells were unable to form large clones in soft agar (Figure S6C, D, F, G, I, J), an indication of inhibited cell proliferation and growth. Then we tested how the tumorigenic potential was affected in the knockdown cells. A375 cells formed tumors when injected subcutaneously into immunodeficient nude mice. However, knockdown of PRMT1 in the cells abolished the tumorigenicity of A375 cells, because no tumor formation by the knockdown cells was observed (Figure 6A-C). The NE-4C cells formed tumors in all injected mice, as we reported recently (Xu et al., 2020), Prmt1 knockdown severely compromised its tumorigenic potential (Figure 6D-F). We compared gene expression representing different cell differentiation between the tumors derived from control NE-4C cells and the tumors derived from the cells with NE-4C knockdown. There was a general tendency that expression of genes representing neural stemness (Figure 6G) and genes representing neuronal differentiation (Figure 6H) was lower in knockdown tumors than control tumors. Moreover, genes denoting endodermal tissue differentiation (*Afp*, *Gata4*, *Sox17*) and epithelial cells (*Cdh1*) was significantly upregulated, whereas the genes for mesodermal tissue differentiation (*Desmin*, *Myh4*, *Acta2*, *Myog*) were downregulated in knockdown tumors (Figure 6I). Moreover, there was a reduction in the expression of the epigenetic factors and the components of the proteasome and ribosome in knockdown tumors (Figure 6J). Our previous work suggested that tumorigenicity and differentiation potential stem from neural stemness (Xu et al., 2020; Cao, 2020). PRMT1/Prmt1 knockdown causes a neuronal differentiation effect and thus loss or reduced neural stemness in cancer cells or NSCs, it is coherent that knockdown cells show a loss of or reduced tumorigenicity. Additionally, tissue or cell differentiation in tumors from NE-4C knockdown cells was compromised as compared with differentiation in tumors from control cells. Knockdown tumors show a differentiation bias towards endodermal tissues but ectodermal (neural) and mesodermal differentiation was both undermined.

**Figure 6.**
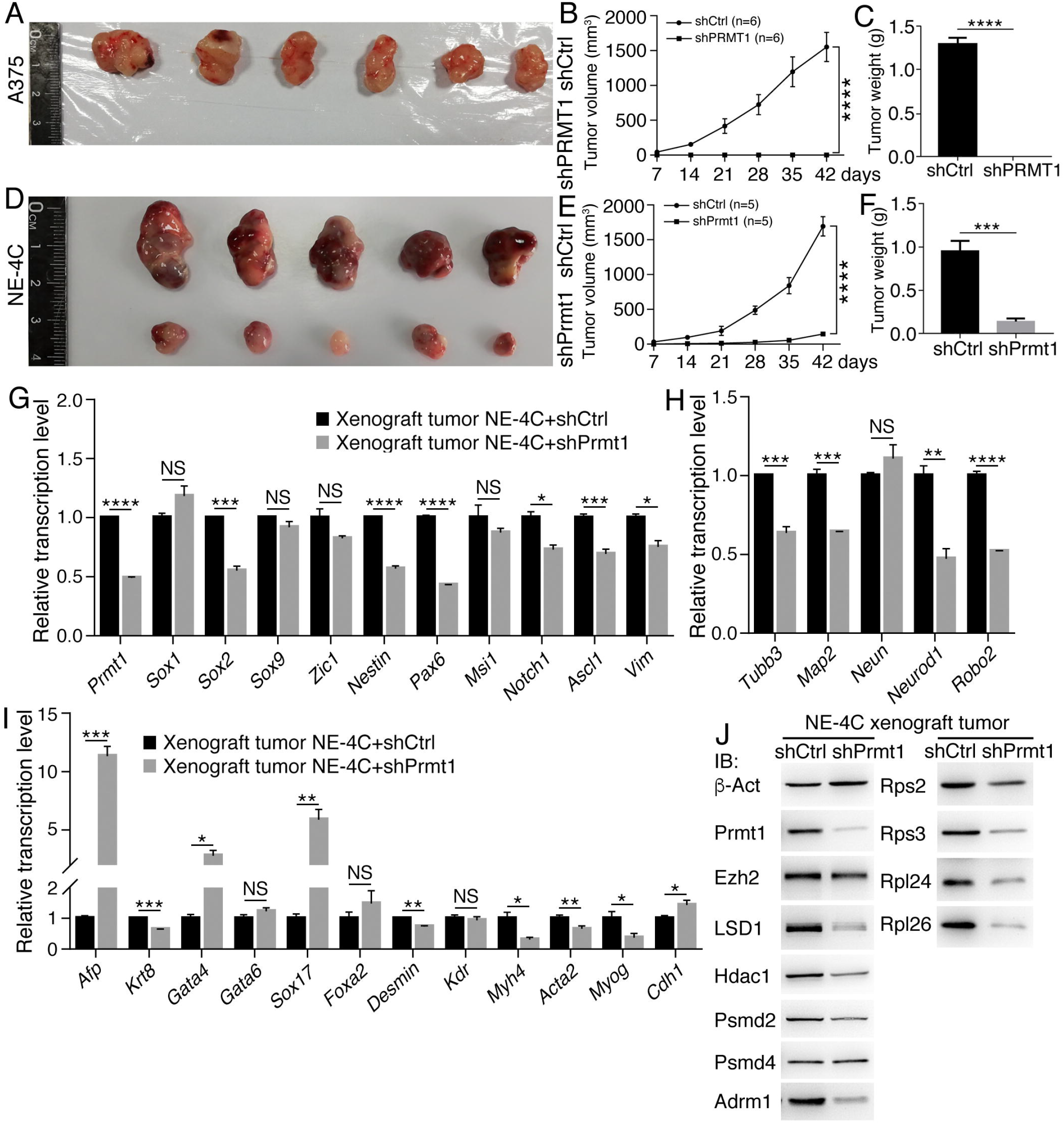
Inhibitory effect of PRMT1/Prmt1 knockdown on cell tumorigenicity and differentiation potential. (A-C) Tumor formation of control (shCtrl) and knockdown (shPRMT1) A375 cells in each six injected nude mice (A), and difference in tumor volume (B) and weight (C) between the two groups. (D-F) Tumor formation of control (shCtrl) and knockdown (shPrmt1) NE-4C cells in each five injected nude mice (D), and difference in tumor volume (E) and weight (F) between the two groups. In (B) and (E), significance of difference in tumor volume between two groups of mice was calculated using two-way ANOVA-Bonferroni/Dunn test. In (C) and (F), significance of difference in tumor weight was calculated using two-tailed Student’s *t*-test. Data are shown as mean±SEM. ***p<0.001, ****p<0.0001. (G-I) Comparison of expression of genes representing neural stemness (G), neuronal differentiation (H), and mesodermal and endodermal differentiation (I) between xenograft tumors derived from control NE-4C cells (NE-4C+shCtrl) and tumors from NE-4C knockdown cells (NE-4C+shPrmt1), as detected with RT-qPCR. Significance in expression change was calculated for experiments in triplicate using two-tailed Student’s *t*-test. Data are shown as mean±SEM. *p<0.05, **p<0.01, ***p<0.001, ****p<0.0001. NS: not significant. (J) Protein expression difference between tumors from control NE-4C cells and tumors from knockdown cells. Pooled protein samples of the control and knockdown groups, respectively, were used for IB.

## Discussion

Based on our studies in recent years and many other pieces of evidence, we propose that neural stemness represents the ground or basal state of cell tumorigenicity and pluripotent differentiation potential (Zhang et al., 2017; Lei et al., 2019; Xu et al., 2020; Cao, 2020). The two cell properties are both derived from the property of neural stemness, which is pre-determined by evolutionary advantage (Xu et al., 2020; Cao et al., 2020). Ribosomes, proteasomes and epigenetic modification factors are usually considered to function in all cell types of an organism and not cell type specific. This is true if the level of their expression or activity is not considered. In fact, the genes for the machineries for basic cellular physiological functions and developmental programs, such as cell cycle, ribosome biogenesis, translation, proteasome assembly, RNA splicing, epigenetic modification, reprogramming, etc., are mostly enriched in embryonic neural cells, as shown in Figure S2 and summarized in Cao (2020). These machineries work together to establish the property of neural stemness, a basal and most proliferative state. A highly proliferative state needs high activity of both ribosomes and proteasomes, and low activity of these machineries fits for a low or non-proliferative state. This needs a coordination to achieve such a balance. The coordination is firstly reflected by that expression of the genes of many epigenetic factors including PRMT1, ribosome and proteasome components, etc., is enriched only in embryonic neural cells, and by that these factors are upregulated in and promote cancers. When either NSCs or cancer cells differentiate into neuronal or neuronal-like cells, either by treatment with RA or inhibition of PRMT1, PRMT1, ribosomal and proteasomal components are downregulated. In the meanwhile, epigenetic factors like EZH2, LSD1 and HDAC1, etc., whose expression is enriched in embryonic neural cells, are reduced. During normal neural development, expression of these proteins is decreased with the progression of neural development (Chau et al, 2018; present study). Differentiation into muscle-like cells causes the same tendency of expression change, which mimics the difference in expression levels of the genes for these proteins between neural and non-neural cells in early embryos (Figure S2; Cao, 2020). It was reported that differentiation of ESCs, whose default fate is primitive neural stem cells (Tropepe et al., 2001; Muñoz-Sanjuán and Brivanlou, 2002; Smukler et al., 2006; Gilbert and Barresi, 2016), is accompanied with downregulation of the proteasome component PSMD11. It is thus deduced that cleanup is necessary for maintaining pluripotency (Schröter and Adjaye, 2014). Our suggestion is that a high level of proteasome activity is required for matching high protein production in a highly proliferative state. By contrast, dedifferentiation by loss of Myod1 in myoblast cells leads to the gain of the property of neural stemness and tumorigenicity (Xu et al., 2020). Coherently, the epigenetic factors and the ribosome and proteasome components are upregulated. Taking together, neural stemness, representing a basal and highly proliferative state, is defined by a high level of the machineries required for basic cell physiological functions. These machineries should be, and actually are, repressed in differentiated cells. Consequently, loss of neural stemness via differentiation leads to loss of or reduced tumorigenicity in cancer cells (Zhang et al., 2017; Xu et al., 2020; present study). Reversal of the differentiated state, like that occurs during tumorigenesis, restores a high level of these machineries again.

It needs a coordination mechanism to achieve a balanced level of different machineries. PRMT1 might play such a role. PRMT1 interacts with many ribosome and proteasome components, places protective marks of methylation of arginine residues on ribosome or proteasome proteins. Decrease in PRMT1 results in losing the protective marks on the proteins, ultimately leading to not only proteasomal degradation of these proteins, but also the alteration of their subcellular distribution. Ribosome and proteasome component proteins tend to distribute primarily in the cytoplasm in response to PRMT1 inhibition. The changes cause the disruption of ribosome biogenesis and proteasome assembly. Besides our observation of PRMT1 regulation of ribosome biogenesis via RPS3 and RPL26 methylation, it seems that ribosome biogenesis is more extensively tuned by different PRMTs, primarily via methylation of different ribosome components (Chern et al., 2002; Bachand and Silver, 2004; Swiercz et al., 2005; Shin et al., Ren et al., 2010). Regulation of proteasome assembly by PRMTs has not been reported. Reduced expression of EZH2, LSD1 and HDAC1 in response to blocking of PRMT1 might be not only due to proteasomal degradation. Disrupted ribosome biogenesis should also play a part in their expression level.

The roles of PRMT1 are not limited to the regulation of ribosome biogenesis and proteasome assembly. Our mass spectrometric assay also identified the components of the machineries for spliceosome and translation initiation, etc., as putative interaction partners of PRMT1, suggesting that it might function more extensively in the regulation of basic cellular physiological processes. In fact, loss of PRMT1 causes decreased arginine methylation of the translation initiation complex, leading to a defect of its assembly and translation and consequently, a repression effect on cancer cells (Hsu et al., 2017). The association between PRMTs, including PRMT1, and RNA splicing was also investigated (Zhang et al., 2015; Fong et al., 2019). Hence, PRMT1 is involved in the coordination of basic cell physiological machineries, which work together to define a basal and highly proliferative property in either NSCs or cancer cells.

## Supporting information

Putative Prmt1 interaction partners in NE-4C cells identified with mass spectrometry

Putative PRMT1 interaction partners in HepG2 cells identified with mass spectrometry

## Author contributions

Y.C. conceived the research. L.C. and M.Z. performed cell and animal experiments; L.C. performed biochemical experiments; L.F. and L.C. performed mass spectrometry; L.C., X.Y., L.X., L.S. and N.C. performed molecular experiments; Y.C. wrote the manuscript.

## Materials and methods

### Cell culture

C57BL/6 mESCs were cultured in Dulbecco’s Modified Eagle’s Medium (DMEM. Thermo Fisher Scientific, #11965-092), 1 ng/ml human LIF (Cell Signaling, #8911), 100 μM β-mercaptoethanol, 2 mM L-glutamine (Thermo Fisher Scientific, #25030164), 1×MEM nonessential amino acids (Thermo Fisher Scientific, #11140050). A375, HepG2, C2C12^*WT*^, and C2C12^*Myod1*−/−^ were cultured in DMEM. A549 was cultured in F-12K medium. NE-4C cells were cultured in MEM (Gibco, #11090073) containing 1% glutamax (Gibco, #35050061) and 1% MEM non-essential amino acids. All culture media were supplemented with 10% fetal bovine serum (FBS. Gibco, #10099141) (15% FBS for mESCs) and with 50 U/ml penicillin and 50 μg/ml streptomycin. Cells were cultured at 37°C with 5% CO_2_. mESCs were cultured in petri dishes coated with 0.1% gelatin or with MEF feeder cells, and NE-4C was cultured in dishes coated with 10 μg/ml poly-D-lysine (PDL. Sigma-Aldrich, #P0899).

HepG2 (#SCSP-510), A375 (#SCSP-533), A549 (#TCHu150) and NE-4C (#SCSP-1501) were purchased from the Cellbank of Chinese Academy of Sciences (Shanghai, China). mESCs were a gift from Dr. Jinzhong Qin at the Model Animal Research Center of the Medical School, Nanjing University. Cancer cell lines were authenticated with short tandem repeat profiling, and cells were detected free of mycoplasma contamination with PCR.

### In vitro neural differentiation of mESCs and chemical treatment of cells

Primitive NSCs (primNSCs) were differentiated from mESCs by culturing mESCs in defined serum-free NSC medium Ndiff 227 (CellArtis, #Y40002) at 37°C with 5% CO_2_. PrimNSCs formed free-floating neurospheres in the medium four days later and the neurospheres grew larger with continuing culture. Wild type or cells with PRMT1/Prmt1 knockdown were also cultured in this medium to test their ability in neurosphere formation and neuronal differentiation effect. To induce neuronal differentiation in NE-4C cells, retinoic acid (RA. Sigma-Aldrich, #R2625) was added to the culture medium to a final concentration of 1 μM for 24 hours. Later on, RA was removed and cells were cultured further till significant differentiation occurred, as indicated in the text. HEK293T cells were treated with MG132 (Selleckchem, #S2619) at 25 μM for 18 hours to detect protein ubiquitination, and A375 cells were treated with MG132 at 25 μM for 18 hours for the detection of protein expression. To examine protein half-life, HEK293T cells that were infected with control virus (shCtrl) or PRMT1 knockdown (shPRMT1) virus were treated with cycloheximide (CHX. Selleckchem, #S7418) at a final concentration of 50 μg/ml in a time series of 0, 2, 4, 6 and 8 hours, respectively.

### Immunofluorescence (IF)

IF detection of protein expression in cells or neurospheres was performed using conventional method as described (Xu et al., 2020). Primary antibodies were EZH2 (Cell Signaling Technology, #5246), HDAC1 (Cell Signaling Technology, #5356), LSD1 (Cell Signaling Technology, #2139), MAP2 (Abcam, #ab183830), MEF2C (Cell Signaling Technology, #5030), MYOD1 (Novus Biologicals, #NB100-56511), NEUROD1 (Cell Signaling Technology, #4373), NF-L (Cell Signaling Technology, #2837), PAX6 (Abcam, #ab195045), PRMT1 (Cell Signaling Technology, #2449), PSMA2 (Abclonal, #A2504), PSMD2 (Abclonal, #A1999), RPL26 (Abclonal, #A16680), RPS3 (Abclonal, #A2533), SOX1 (Abcam, #ab109290), TUBB3 (Cell Signaling Technology, #5568), 20S α+β (Abcam, #ab22673). Secondary antibodies were anti-mouse IgG (FITC-conjugated) (Sigma-Aldrich, #F9137), anti-mouse or rabbit Alexa Flour 594 (ThermoFisher Scientific, #A21207, #A21203). Cell nuclei were counterstained with DAPI. After staining, slides were washed and mounted with SlowFade™ Gold Antifade Mountant (ThermoFisher, #S36936). Cells were viewed and photographed with a fluorescence microscope (Zeiss LSM 880).

### Immmunoblotting (IB)

Whole cell lysates (WCL) were prepared and IB detection of protein expression in cells was carried out using conventional methods. Blots were developed with a Western blotting substrate (Tanon, #180-501). Primary antibodies were: β-Act (Cell Signaling Technology, #4970), ADRM1 (Abclonal, #A4481), AURKA (Cell Signaling Technology, #14475), DNMT1 (Abcam, #ab13537), EZH2 (Cell Signaling Technology, #5246), HDAC1 (Cell Signaling Technology, #5356), LSD1 (Cell Signaling Technology, #2139), MAP2 (Abcam, #ab183830), MEF2C (Cell Signaling Technology, #5030), MSI1 (BioLegend, #869101), MYC (Abcam, #ab32072), MYOD1 (Novus Biologicals, #NB100-56511), NEUROD1 (Cell Signaling Technology, #4373), NF-L (Cell Signaling Technology, #2837), PAX6 (Abcam, #ab195045), PCNA (Cell Signaling Technology, #13110), PRMT1 (Cell Signaling Technology, #2449), PSMA2 (Abclonal, #A2504), PSMD2 (Abclonal, #A1999), PSMD4 (Abcam, #ab137109), RPL24 (Abclonal, #A14255), RPL26 (Abclonal, #A16680), RPS2 (Abclonal, #A6728), RPS3 (Abclonal, #A2533), SETDB1 (Abcam, #ab107225), SOX1 (Abcam, #ab109290), SOX2 (Cell Signaling Technology, #23064), SYN1 (Cell Signaling Technology, #5297), TUBB3 (Cell Signaling Technology, #5568), and ZIC1 (Abcam, #ab134951). The secondary antibodies were HRP-conjugated goat anti-rabbit or anti-mouse IgG (Sangon Biotech, #D110058; #D110087).

### Plasmid construction, viral production, cell infection or transfection

For functional knockdown of PRMT1/Prmt1 and MDN1, a shRNA-based approach was used. The sequences of shRNAs against human PRMT1, MDN1 and mouse Prmt1 were the validated MISSION shRNAs (Sigma-Aldrich). The shRNAs were TRCN0000290479 (human *PRMT1*), TRCN0000018491 (mouse *Prmt1*) and TRCN0000229952 (human *MDN1*), and subcloned to the lentiviral vector pLKO.1. For transient overexpression of human RPS3, RPL26 and PSMD2, the open reading frames (ORFs) were subcloned to pCS2+4×HAmcs or pCS2+6×MTmcs vectors. For overexpression of PRMT1 or MYOD1, the ORFs was subcloned to pLVX-IRES-puro and pLVX-IRES-ZsGreen lentiviral vectors. PRMT1 ORF was also subcloned to pCS2+6×MTmcs vector for the use for transient overexpression.

Viral production and cell infection were performed essentially as described (Lei et al., 2019). For stable knockdown or overexpression assays, virus packaging plasmids and shRNA or overexpression constructs were transfected into HEK293T cells using polyethylenimine (PEI). Polybrene at a final concentration of 10 μg/ml was added to the lentiviral supernatant 48 hours after transfection. The supernatant was centrifuged at 4°C to concentrate the lentiviral particles, which were used for infecting cells. Forty-eight hours after infection, cells were selected with puromycin at 2 μg/ml in culture for 3 days when a puromycin selection vector was used. Cells were cultured further until a desired time when a significant phenotypic change was observed or they were harvested for additional assays. Virus packaging with the empty vector and cell infection with the virus were performed in parallel, which was used as a negative control for knockdown or overexpression assay, respectively.

For transient overexpression assays, HEK293T cells, either untreated or after infected with virus for knockdown of PRMT1 or MDN1 or empty vector control, were transfected with overexpression plasmid using PEI when cells grew to 70-80% confluency. Fourty-eight hours later, cells were collected for biochemical assays.

### Co-IP, mass-spectrometry and functional annotation analysis

Co-IP was performed using a conventional method as described (Lei et al., 2019). Cells were harvested, washed twice with cold PBS and resuspended in lysis buffer (300 mM NaCl, 1% NP-40, 2 mM EDTA, 50 mM Tris-Cl pH7.5 and protease inhibitor cocktail). After 20 min on ice, cells were centrifuged for 20 min at 12000 rpm. The supernatant was collected for immunoprecipitation using the antibody against IgG, an endogenous protein, or an HA- or MT-tag, which was linked to protein G sepharose beads. After incubation overnight at 4°C, beads were washed in TBST buffer (25 mM Tris-Cl PH7.2, 150 mM NaCl, 0.5% Tween-20). Immuno-complexes were eluted by incubating the beads in 1× loading buffer at 95°C, and assayed with SDS-PAGE.

To identify PRMT1 interaction proteins with mass spectrometry, Co-IP was carried out in the same way above using NE-4C and HepG2 cells and an anti-PRMT1/Prmt1 antibody (Cell Signaling Technology, #2449). Precipitated proteins were subjected to SDS-PAGE, which was run for a short time to allow proteins to be stacked and migrate into separating gel. The gel slice containing all proteins was excised and processed. For in gel protein digestion, cysteine residues were reduced by addition of 25 mM final of dithiothreitol (DTT) for 60 min at 70°C, and alkylated by addition of iodoacetamide at a final concentration of 90 mM for 30 min at room temperature in the dark. The proteins were then digested overnight at 37°C with 0.2 μg of modified sequencing grade trypsin (Promega) in 50 mM ammonium bicarbonate. The resulting peptides were extracted from the gel by incubation in 50 mM ammonium bicarbonate: acetonitrile (1:1) for three times of 15 min at 37°C, and then dried and resuspended with 10 μl of 3% acetonitrile, 2% FA before being subjected to LC-MS/MS analysis. Data acquisition was performed with a Triple TOF 5600 System (AB SCIEX, Concord, ON). For database searching, the original files were submitted to ProteinPilot Software (version 4.5, AB Sciex) for data analysis. LC-MS/MS data were searched against *Mus musculus* UniProt database (April 9, 2016, containing 50,943 sequences, http://www.uniprot.org/proteomes/UP000000589) or *Homo sapiens* UniProt database (April 9, 2016, containing 160,566 sequences, http://www.uniprot.org/proteomes/UP000005640) using the parameters: TripleTOF 5600, trypsin digestion and thorough ID. All other parameters were set as default in ProteinPilot Software. A protein is considered to be a putative PRMT1/Prmt1 interaction partner when a protein peptide(s) are identified in the immunoprecipitate with PRMT1/Prmt1 antibody but not in the precipitate with IgG antibody, or the fold change between the number of a protein peptide(s) in the sample precipitated by PRMT1/Prmt1 antibody and by IgG antibody is ≥ 3 (Tables S1 and S2).

Functional annotation for the genes of the putative PRMT1/Prmt1 binding proteins was performed using the DAVID annotation tools (Huang et al., 2007) with default settings.

### Total RNA preparation and reverse transcriptase-quantitative polymerase chain reaction (RT-qPCR)

Total RNAs were extracted from cells or tumors with TRIzol. cDNAs were transcribed from the total RNAs with the HiScript II 1st Strand cDNA Synthesis Kit (+gDNA wiper) (Vazyme, #R212-01/02), which contains the reagent for cleaning up the contamination of genomic DNA. qPCR was performed on a LightCycler® 96 system (Roche) using the following parameters: one cycle of pre-denaturation at 95°C for 5 min, followed by 40 cycles of denaturation at 95°C for 10 sec, annealing and extension at 60°C for 30 sec, and an additional cycle for melting curve. β-Act/β-ACT was detected as a loading control. Significance of changes in transcription was calculated based on experiments in triplicate using two-tailed Student’s *t*-test. Data were presented as histograms with relative units of transcription levels. Primers for RT-qPCR are listed in Table S3.

### Gene expression profiling assay on A549 cells with knockdown of PRMT1

A549 cells were infected with lentivirus carrying the empty vector (shCtrl) or carrying shPRMT1, respectively, and selected with puromycin. After significant phenotypic change occurred in cells with infection of shPRMT1 virus, both control and knockdown cells were collected and subjected to gene expression microarray. RNA extraction/purification, cRNA probe synthesis, probe hybridization to microarrays, signal processing, raw data analysis, and the subsequent enrichment and annotation of pathway and gene ontology were as exactly described (Zhang et al., 2017). Results are shown as bar charts. Raw data are deposited in GEO under accession number GSE162840.

### Mouse embryonic cortical cell isolation

All animal use in the research was approved by and in accordance with the guidelines of the Institutional Animal Care and Use Committee (IACUC) at the Model Animal Research Center of the Medical School, Nanjing University. For isolation of embryonic cortical cells, mouse embryos at E13.5 and E15.5 were resected and brains were excised from embryos. Cortical tissues were separated after removal of meninges and the ganglionic eminences. Cortical tissues were washed and homogenized in cold RIPA lysis buffer (20 mM Tris-HCl, pH 7.4, 150 mM NaCl, 1 mM EDTA, 1% NP-40, 0.5% sodium deoxycholate, and 0.1% SDS) supplemented with protease inhibitors (Roche, #04693132001). Lysates were cleared by centrifugation (14,000 rpm for 15 min) and subjected to immunoblotting.

### Cell migration/invasion assays

Cell migration assay was performed in 24-well transwell plates with inserts of 8-μm pore size (Corning, #3422). Each 1×10^5^ cells were suspended in 200 μl of serum-free culture medium, and added to the upper compartment. 500 μl of culture medium containing 10% FBS was added in the lower compartment. Plates were incubated at 37 °C for a desired time period, as indicated in the text. Afterwards, cells were fixed with 37% formaldehyde, stained with 0.5% crystal violet for 10 min. Cells that did not migrate were removed. Migrated cells were washed with PBS and photographed.

For cell invasion assay, each 80 μl of Matrigel (Corning, #354234) was diluted in PBS (1:8) and distributed uniformly onto a 24-well transwell insert. 2×10^5^ A549 or NE-4C cells or 5×10^5^ A375 cells were added to Matrigel. Plates were incubated at 37 °C for desired time periods as indicated in the text. Afterwards, cells were processed in the same way as in the migration assay.

### Soft agar colony formation assay

The top and bottom layer of agar was 0.35% and 0.7%, respectively, of low melting agarose (BBI, #AB0015). Each 3000 cells were distributed in a well of a 6-well culture plate and cultured for desired time periods, as indicated in the text. Each experiment was performed in triplicate. Significance of difference in colony formation was calculated using two-tailed Student’s *t*-test.

### Xenograft tumor assay

Immunodeficient nude Foxn1^nu^ mice with an age of five to six weeks were purchased from the National Resource Center for Mutant Mice (Nanjing, China) and maintained in a specific-pathogen-free facility. In order to examine the change in tumorigenicity of A375 or NE-4C cells in response to PRMT1/Prmt1 knockdown, cells infected with control virus or with knockdown virus were selected with puromycin. 3×10^6^ A375 or 1×10^6^ NE-4C cells, either control or knockdown, were suspended in 100 μl of sterile PBS, and injected subcutaneously into the dorsal flank of a mouse, respectively. Tumor growth was measured periodically, as indicated in the text. At the end of tumor growth, mice were sacrificed. Tumor tissues were excised and weighed. Tumor volume was calculated using the formula: length×width^2^/2. The significance of difference in tumor volumes between control and knockdown groups was calculated using two-way ANOVA followed by Bonferroni/Dunn (ANOVA-Bonferroni/Dunn) tests. Significance of difference in tumor weight between control and knockdown groups were calculated with two-tailed Student’s *t*-test.

