## Supplemental Figures S1-S6 and figure legends, and supplemental Table S3 and table legends S1-S3 for "Coordinated regulation of ribosomes and proteasomes by PRMT1 in the maintenance of neural stemness of cancer cells and neural stem cells"

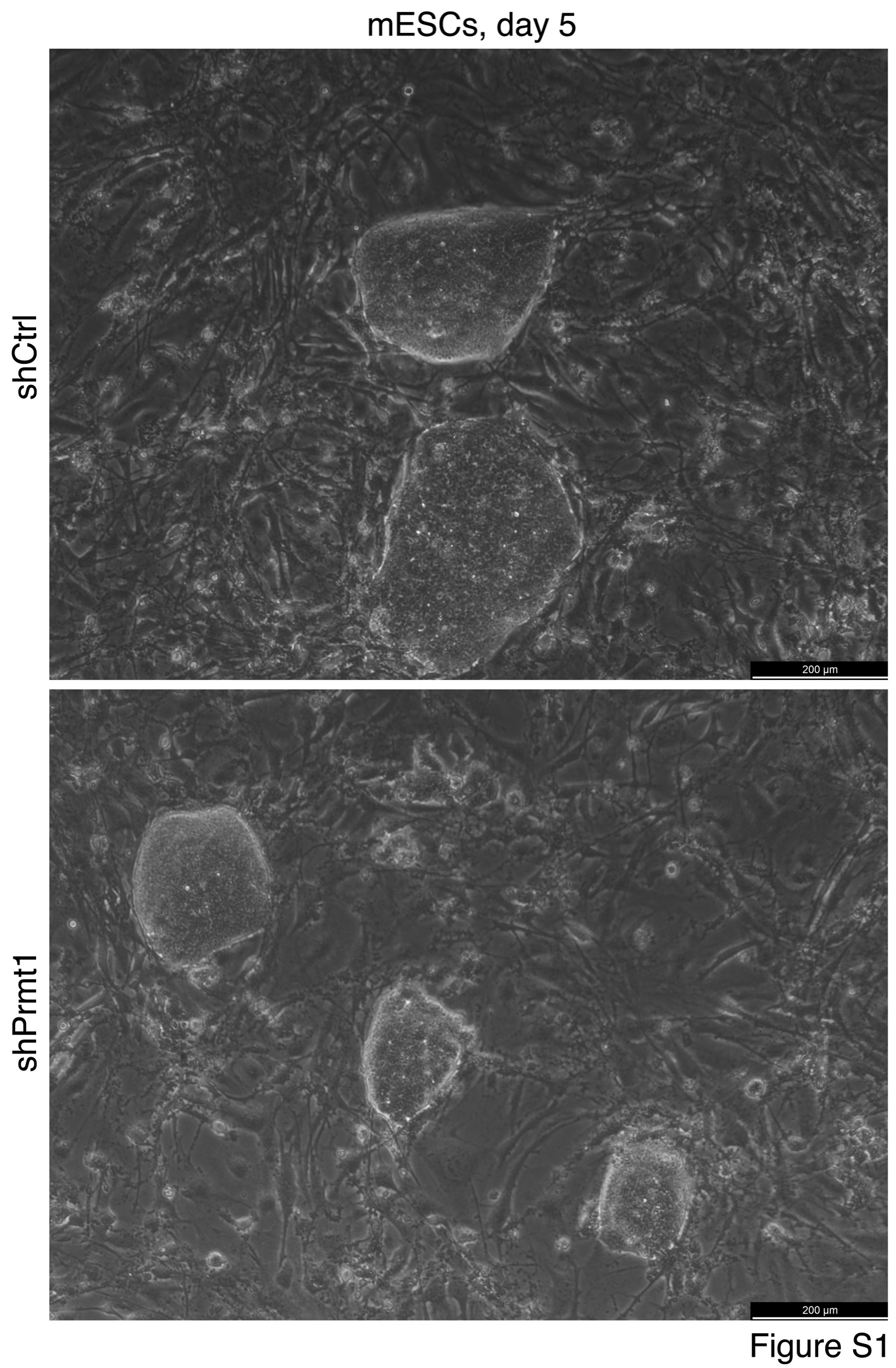

**Figure S1. Colony formation of control mESCs (shCtrl) and mESCs with Prmt1 knockdown (shPrmt1). Related to Figure 1.**

Representative colonies were photographed five days after cells were seeded on feeder cell layers.

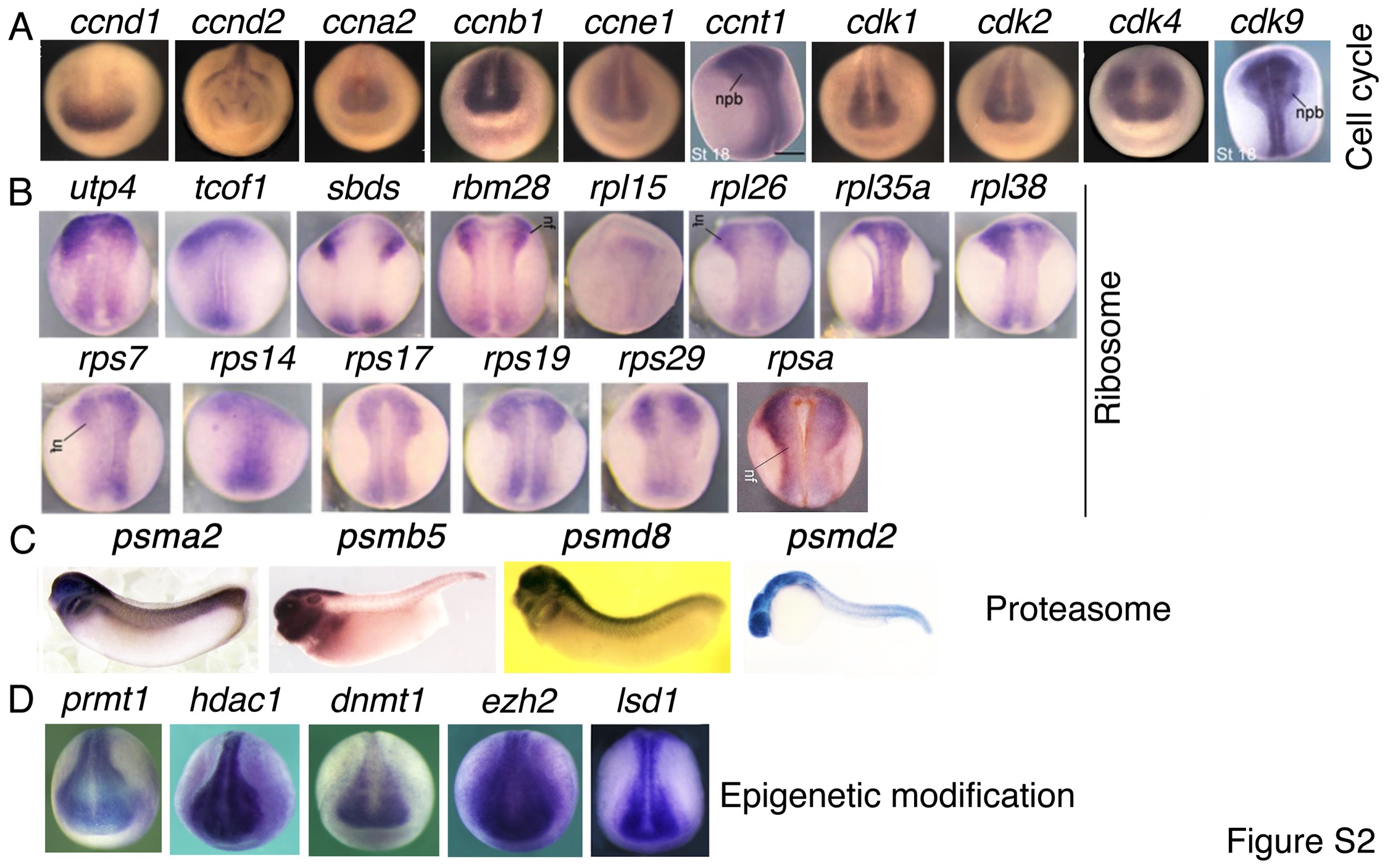

**Figure S2. Neural enriched expression of genes for machineries for basic cell physiological functions during vertebrate embryogenesis. Related to Figure 3.**

(A) Cell cycle. (B) Ribosome biogenesis. (C) Proteasome assembly. (D) Epigenetic factors. Shown here are expression patterns of genes in *Xenopus* except *psmd2*, for which the expression pattern in zebrafish is displayed since its expression in *Xenopus* embryos is not known. *Xenopus* embryos are at neurula stages (A, B, D) and tailbud stages (C), and the zebrafish embryo is at stage Prim-5. Data of gene expression patterns were from the *Xenopus* ([www.xenbase.org](http://www.xenbase.org)) or zebrafish ([www.zfin.org](http://www.zfin.org)) databases, and adapted from Cao (2020).

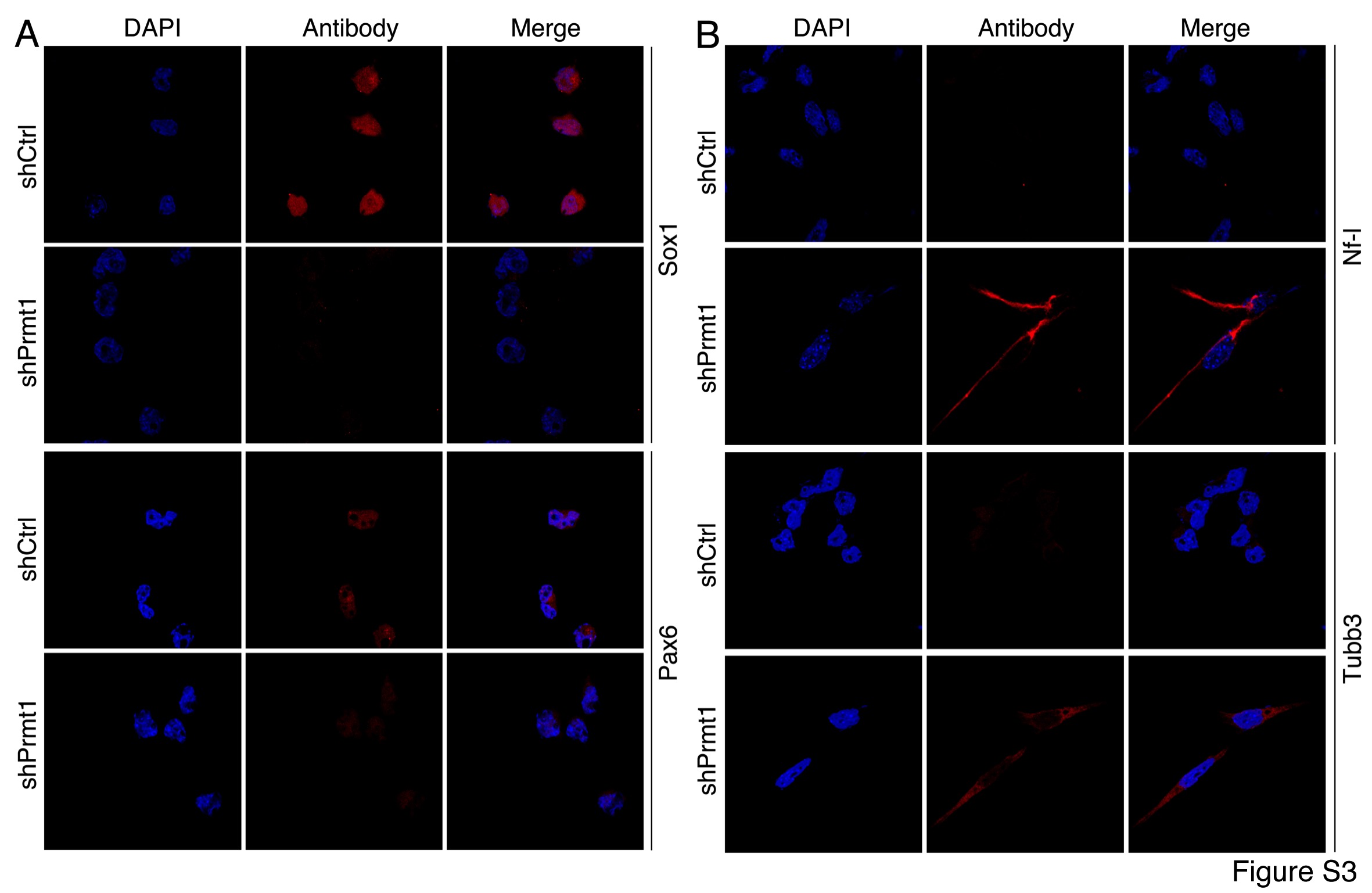

**Figure S3. Detection of neuronal differentiation of NE-4C cells after knockdown of Prmt1. Related to Figure 3.**

(A) Neural stemness marker expression in control and knockdown cells detected with IF. (B) Neuronal marker expression in control and knockdown cells. Nuclei were counterstained with DAPI.

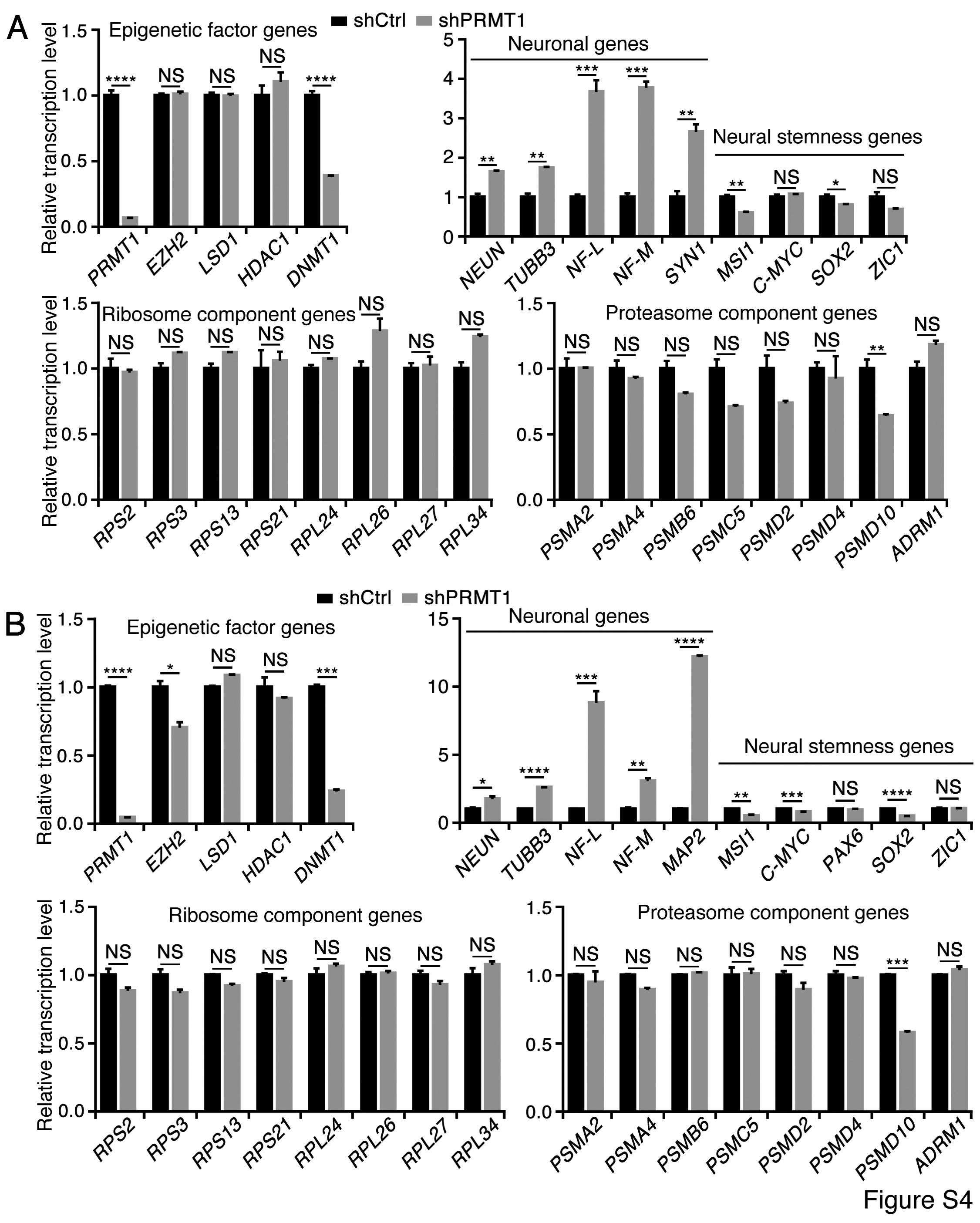

**Figure S4. Gene expression analysis in A375 (A) and A549 (B) cells in response to PRMT1 knockdown using RT-qPCR. Related to Figure 5.**

Significance of gene expression change was calculated for experiments in triplicate using two-tailed Student’s *t*-test. Data are shown as mean±SEM. *p<0.05, **p<0.01, ***p<0.001, ****p<0.0001. NS: not significant.

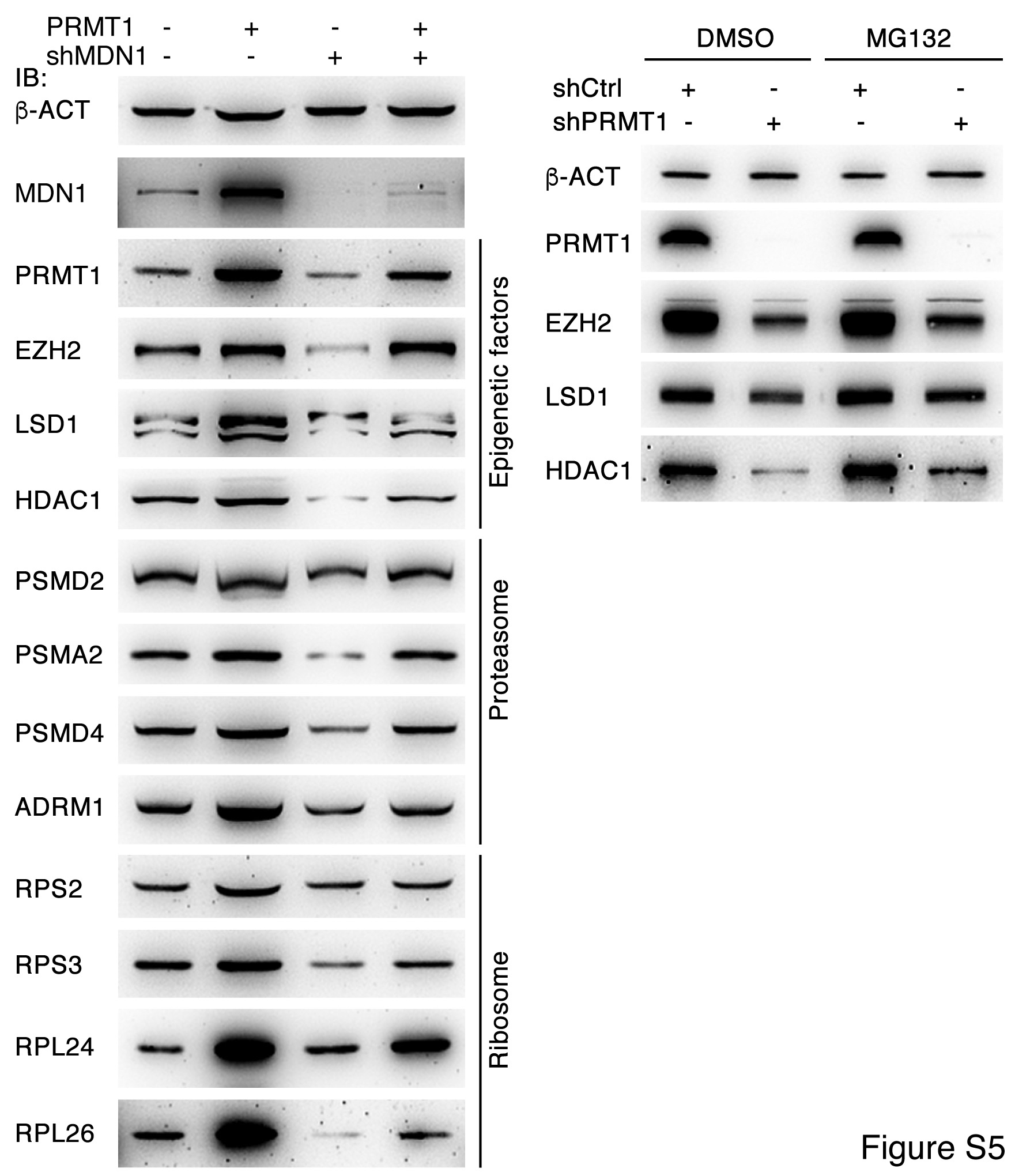

**Figure S5. The Effect of inhibition of ribosome biogenesis or proteasome activity on protein expression. Related to Figure 5.**

(A) Inhibition of ribosome biogenesis via knockdown of MDN1 in A549 cells led to downregulation of protein expression, which can be reversed by overexpression of PRMT1. (B) The effect of inhibition of proteasome activity via treating A375 cells with MG132 can be weakened by knockdown of PRMT1.

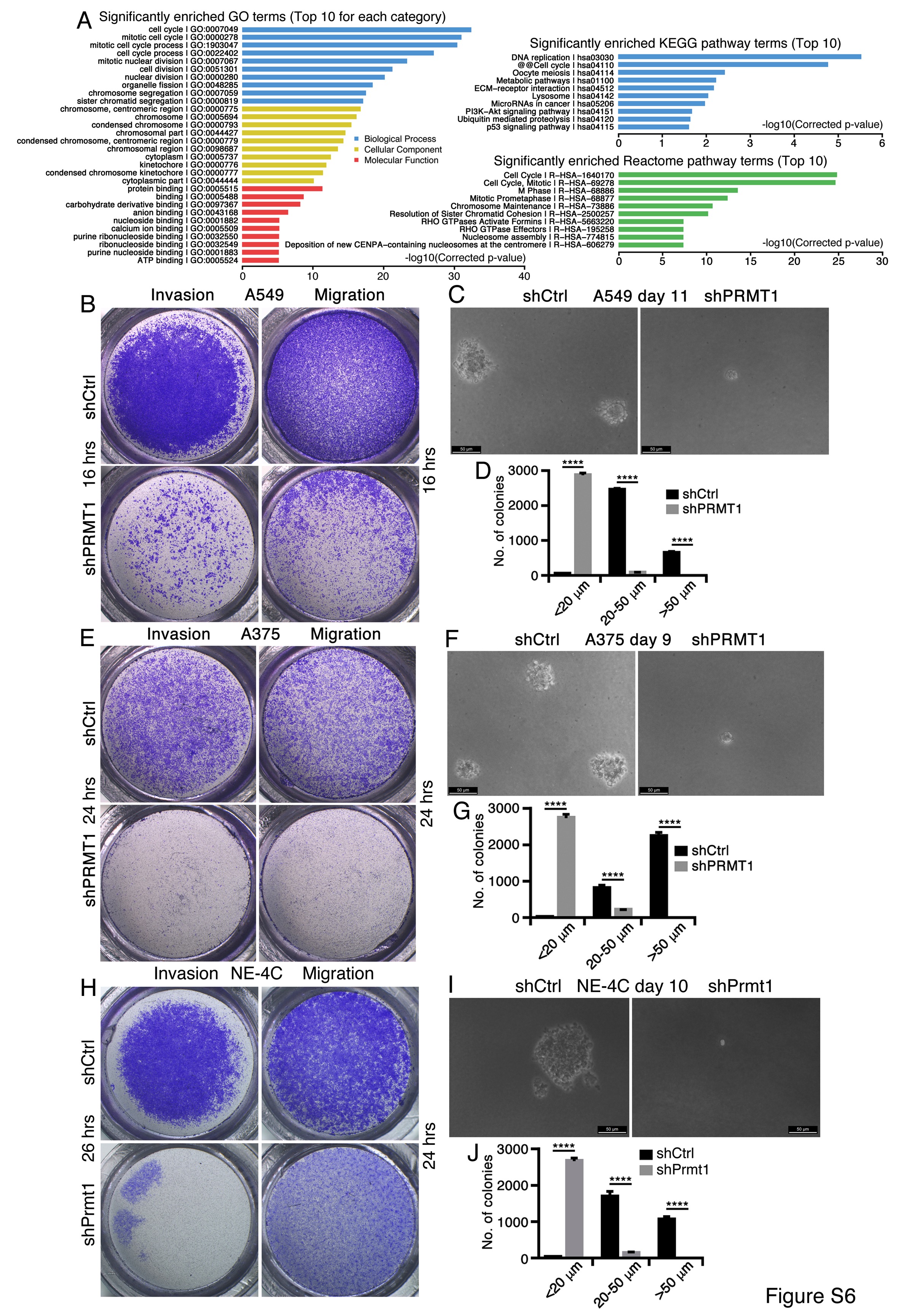

**Figure S6. PRMT1 knockdown alters transcriptome and malignant features in cancer or NSC cells. Related to Figure 6.**

(A) Enriched GO and pathway terms for the altered genes in A549 cells with knockdown of PRMT1. (B-J) Cancer cells A549 (B-D) and A375 (E-G) and the neural stem cell NE-4C (H-J) show repressed capability of invasion and migration (B, E, H), and colony formation in soft agar (C, F, I), in response to PRMT1/Prmt1 knockdown. (D), (G) and (J) show the significance of difference in colony formation in control and knockdown cells based on experiments in triplicate using two-tailed Student’s *t*-test. Data are shown as mean±SEM. ****p<0.0001.

**Table S1. Putative Prmt1 interaction partners in NE-4C cells identified with mass spectrometry. Related to Figure 2.**

**Table S2. Putative PRMT1 interaction partners in HepG2 cells identified with mass spectrometry. Related to Figure 2.**

**Table S3. Primers for RT-qPCR. Related to Figures 5 and 6.**

| Mouse genes | Primers (5’-3’) |
| --- | --- |
| *Acta2* (*alpha-Sma*) | Forward: aatggctctgggctctgtaa Reverse: tctcttgctctgggcttcat |
| *eta-Act* | Forward: ccctgaagtaccccattgaa Reverse: cttttcacggttggccttag |
| *Afp* | Forward: atgaagcaagccctgtgaac Reverse: agcttggcacagatccttgt |
| *Ascl1* | Forward: gccaacaagaagatgagcaag Reverse: gaacccgccatagagttcaa |
| *Cdh1* | Forward: actttggtgtgggtcaggaa Reverse: ttcacatgctcagcgtcttc |
| *Desmin* | Forward: gtgaagatggccttggatgt Reverse: cgggtctcaatggtcttgat |
| *Foxa2* | Forward: taagcgagctaaagggagca Reverse: gtggttgaaggcgtaatggt |
| *Gata4* | Forward: ggaagcccaagaacctgaat Reverse: tgctgtgcccatagtgagat |
| *Gata6* | Forward: aagatgaatggcctcagcag Reverse: catatagagcccgcaagcat |
| *Kdr* | Forward: agctctccgtggatctgaaa Reverse: agatgctccaaggtcaggaa |
| *Krt8* | Forward: tctgggatgcagaacatgag Reverse: tcttcacaaccacagccttg |
| *Map2* | Forward: agcagccgaagaaacagcta Reverse: aaggtcttgggagggaagaa |
| *Myh4* | Forward: acattattggctggctggac Reverse: acccttcttcttgccacctt |
| *Myog* | Forward: tccagtacattgagcgccta Reverse: caaatgatctcctgggttgg |
| *Nestin* | Forward: ttccctgatgatccaacctc Reverse: acctctgtggctgcttcttt |
| *Neurod1* | Forward: ctctggagcccttctttgaa Reverse: tgcagggtagtgcatggtaa |
| *NeuN* | Forward: agcagcccaaacgactacat Reverse: tcggtcagcatctgagctagt |
| *Notch1* | Forward: tcgtgctcctgttctttgtg Reverse: agcaccatctgaggcattct |
| *Pax6* | Forward: cacatcaggttccatgttgg Reverse: cataactccgcccattcact |
| *Prmt1* | Forward: gtggatgggttactgcctct Reverse: aagccatacacgttctccca |
| *Robo2* | Forward: attccgttgtcaggtccaag Reverse: acttttcccacccgattctc |
| *Sox1* | Forward: cacaactcggagatcagcaa Reverse: tccttcttgagcagcgtctt |
| *Sox2* | Forward: gcggagtggaaacttttgtc Reverse: tccgggaagcgtgtacttat |
| *Sox9* | Forward: gctggcaaagttgatctgaag Reverse: gttgggtggcaagtattggt |
| *Sox17* | Forward: taaaggtgaaaggcgaggtg Reverse: tagctctgcgttgtgcagat |
| *Tubb3* | Forward: ttctggtggacttggaacct Reverse: actctttccgcacgacatct |
| *Vim* | Forward: gaccttgaacggaaagtgga Reverse: agccacgctttcatactgct |
| *Zic1* | Forward: tggagccttcttccgctat Reverse: actcctcccagaagcagatgt |
| Human genes | Primers (5’-3’) |
| *eta-ACT* | Forward: agaaaatctggcaccacacc Reverse: tagcacagcctggatagcaa |
| *DRM1* | Forward: tacattcagcagacggacga Reverse: cctgcatccagaagaaaagc |
| *NMT1* | Forward: gagccacagatgctgacaaa Reverse: tgccattaacaccaccttca |
| *EZH2* | Forward: ccgctgaggatgtggatact Reverse: cttggtgttgcactgtgctt |
| *HDAC1* | Forward: atatcgtcttggccatcctg Reverse: ggcttgaaaatggcctcata |
| *LSD1* | Forward: atctgcagtccaaaggatgg Reverse: gccaacaatcacatcgtcac |
| *NEUN* | Forward: gagaagctgaatgggacgat Reverse: aaccccgtcactgcatagaa |
| *MAP2* | Forward: cagggaggaatttgtggaga Reverse: atggtctcctttccacctca |
| *MSI1* | Forward: accaagagatccaggggttt Reverse: tcgttcgagtcaccatcttg |
| *NF-L* | Forward: agaccctggaaatcgaagca  Reverse: tcgccttccaagagtttcct |
| *NF-M* | Forward: aaatggaagaggccctgaca  Reverse: tcttcggcttggtctgactt |
| *PAX6* | Forward: aagggccaaatggagaagag Reverse: gccagatgtgaaggaggaaa |
| *PRMT1* | Forward: taccgtcaaggtggaagacc Reverse: gtccaggtcgatggtgaagt |
| *PSMA2* | Forward: tttggtgtacagtggcatgg Reverse: actccaaatggacgaacacc |
| *PSMA4* | Forward: atgaggacatggcttgcagt Reverse: accaaagggacgttttcctc |
| *PSMB6* | Forward: gccaatcgagtgactgacaa Reverse: tcccggtatcggtaacacat |
| *PSMC5* | Forward: aagtgatcgagctgcctgtt Reverse: ttcagagccagagacacgaa |
| *PSMD2* | Forward: agaacaaggaccacggaatg Reverse: cactatgccacaggcaagaa |
| *PSMD4* | Forward: actggctaaacgcctcaaga Reverse: tgagagcatcagccaaactg |
| *PSMD10* | Forward: agcagccaagggtaacttga Reverse: cacttgcaggggtgtctttt |
| *RPL24* | Forward: atgcgagtcggctttccttt Reverse: agagatgcaccagtaatggcc |
| *RPL26* | Forward: tacgtgatctacatcgagcgc Reverse: tctcctccttgtacttgccct |
| *RPL27* | Forward: cattgatgatggcacctcag Reverse: ccttgtgggcattaggtgat |
| *RPL34* | Forward: caagaaggttgggaaagcac Reverse: gaaagcacgcttgatcctgt |
| *RPS2* | Forward: tgaagatcaagtccctggagg Reverse: aaatgccttgaacctggtgc |
| *RPS3* | Forward: atgctgaaaaggtggccact Reverse: tttcccagacaccacaacct |
| *RPS13* | Forward: acgacgtgaaggagcagatt Reverse: aggaagatcaggagcaagtcc |
| *RPS21* | Forward: gaaatgctccgctagcaatc Reverse: tgactcacccatcctacgaa |
| *SOX2* | Forward: catcacccacagcaaatgac Reverse: cctccccaggttttctctgta |
| *SYN1* | Forward: aatactggctctgcgatgct Reverse: tgtcttcatcctggtggtca |
| *TUBB3* | Forward: gtgcggaaggagtgtgaaa  Reverse: acgacgctgaaggtgttcat |
| *ZIC1* | Forward: acaaaaggacgcacacagg Reverse: ggattcgtggaccttcatgt |
